# Learning Latent Trajectories in Developmental Time Series with Hidden-Markov Optimal Transport

**DOI:** 10.1101/2025.02.14.638351

**Authors:** Peter Halmos, Julian Gold, Xinhao Liu, Benjamin J. Raphael

## Abstract

Deriving the sequence of transitions between cell types, or differentiation events, that occur during organismal development is one of the fundamental challenges in developmental biology. Single-cell and spatial sequencing of samples from different developmental timepoints provide data to investigate differentiation but inferring a sequence of differentiation events requires: (1) finding trajectories, or ancestor:descendant relationships, between cells from consecutive timepoints; (2) coarse-graining these trajectories into a *differentiation map*, or collection of transitions between *cell types*, rather than individual cells. We introduce Hidden-Markov Optimal Transport (`HM` -`OT`), an algorithm that simultaneously groups cells into cell types and learns transitions between these cell types from developmental transcriptomics time series. `HM` -`OT` uses low-rank optimal transport to simultaneously align samples in a time series and learn a sequence of clusterings and a differentiation map with minimal total transport cost. We assume that the law governing cell-type trajectories is characterized by the joint law on consecutive time points, tantamount to a Markov assumption on these latent trajectories. `HM` -`OT` can learn these clusterings in a fully unsupervised manner or can generate the least-cost cell type differentiation map consistent with a given set of cell type labels. We validate the unsupervised clusters and cell type differentiation map output by `HM` -`OT` on a Stereo-seq dataset of zebrafish development, and we demonstrate the scalability of `HM` -`OT` to a massive Stereo-seq dataset of mouse embryonic development.

**Code availability:** Software is available at https://github.com/raphael-group/HM-OT

## 1 Introduction

One of the most fundamental questions in biology is how a single zygote cell differentiates into the plethora of functionally specialized and distinct cell types, tissues, and organs which comprise a eukaryotic organism. In the past decades, a great deal of experimental work has elucidated the underlying mechanisms of development, a remarkably conserved process across all animals. From drosophila to humans, development is driven by cascading and spatially-segregated waves of epigenetic and transcriptomic changes. These changes are often summarized in a Waddington landscape [38] which describes cell types or cell states as a location on a landscape (often illustrated in 2D) and with each cell having a developmental potential, or height, describing its differentiation status. Multipotent progenitor cells have high developmental potential and roll down the landscape into differentiated cell types with lower developmental potential (Fig. 1). The key challenge in developmental biology can thus be phrased as learning the Waddington landscape of an organism including (1) the cell types and cell states; (2) a cell differentiation map that describes the developmental trajectories of cells on Waddington’s landscape.

**Figure 1:**
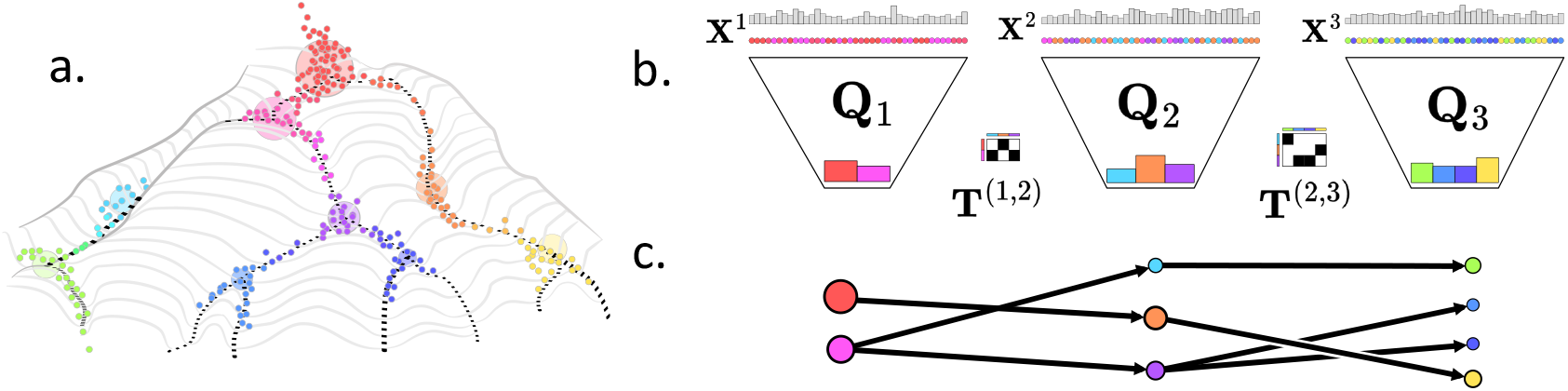
(**a**) Cells (colored points) occupy cell states on a Waddington landscape with undifferentiated cells having high differentiation potential at the top and fully differentiated cells with low differentiation potental at the bottom. Cells cluster into a small number of cell types (large circles). (**b**) Latent representation **Q**_*t*_ is a joint distribution between cells **X**^1^, **X**^2^, **X**^3^ at times 1, 2, 3, (top) and their cell types (bottom). Couplings **T**^(*t,t*+1)^ describe cell type transitions across time. (**c**) A differentiation map summarizes the inferred latent trajectories with vertices indicating cell types and edges indicating transitions.

Over the past few years, many researchers have attempted to derive developmental trajectories and differentiation maps from single-cell or spatial RNA sequencing data obtained from multiple time points during a developmental process. Since these technologies are destructive (an individual cell can be measured only once), the major problem in using this data is to link cells from these multiple time points into trajectories. This problem is usually termed *trajectory inference* and dozens of methods have been developed to solve this problem, with many of the earlier methods benchmarked in a DREAM challenge [28]. A key assumption in most of these methods is that the majority of cells occupy distinct cell types. These methods place cells on a small number of trajectories that connect these cell types, computing a continuous pseudotime for a cell along the trajectory. Other methods use heuristics that examine cells and cell types at consecutive time points. For example, [25, 26] construct cell differentiation maps by heuristics by connecting pairs of cell types from consecutive time points that have high proportions of nearest neighbors in a *k*-NN graph built from the cells at these time points.

Another popular approach for learning cell differentiation maps is to use the framework of *optimal transport* (OT) to learn couplings between two probability distributions over cell states. This approach was pioneered in Waddington-OT [33], a landmark paper and state of the art for single-cell alignment and trajectory inference. More recently, OT has extended to spatial transcriptomics data from multiple time points to learn trajectories in space and time [10, 17, 37]. These and other OT methods learn couplings, or alignments between pairs of *cells* from consecutive time points. Thus, deriving a differentiation map requires post-processing of the inferred couplings, with no guarantees that this coarse-graining is optimal or consistent across multiple time points.

We introduce *Hidden-Markov Optimal Transport (*`HM`*-*`OT`), an algorithm which takes as input a time-series of consecutive single-cell or spatial samples and uses low-rank OT [8, 30, 11] to infer both cell types at each timepoint *and* derive cell type transitions across timepoints. `HM`-`OT` leverages *Factor Relaxation with Latent Coupling (FRLC, mnemonic: “frolic”)* [11] – a new general-purpose algorithm for low-rank OT. FRLC solves for low-rank transport plans factored into three matrices: a pair of *latent representations* (**Q**_*t*_, **Q**_*t*+1_) and a *latent coupling (LC)* matrix (**T**^(*t,t*+1)^) that links the two latent representations. `HM`-`OT` generalizes FRLC to a time-series of any length to derive consistent latent cell type representations (clusterings) across the whole time-series, overcoming limitations of pairwise methods that yield discordant cell type clusterings between time points. `HM`-`OT` optimizes a cost defined on trajectories through the time series which obeys an optimal sub-structure property. Owing to difficulties imposed by continuous dynamic programming, `HM`-`OT` approximates the MAP estimate instead, combining past and future information to learn latent representations and inferring couplings (cluster transitions) linking them into a sequence. The components returned by `HM`-`OT` define a joint distribution on the full time series. This joint distribution is coupled to a Markov chain on the latent (cell type) trajectories through the latent representations (cell type assignments). `HM`-`OT` is also flexible enough to learn a differentiation map *given* cell type annotations at each timepoint; we refer to this as “supervised `HM`-`OT`.”

We demonstrate the advantages of `HM`-`OT` on synthetic and real datasets, including a spatial RNA sequencing dataset of zebrafish embryogenesis from [21], and a spatial RNA-sequencing dataset of mouse embryogenesis [3]. Using information-theoretic metrics, we find that both `HM`-`OT` and supervised-`HM`-`OT` learn differentiation maps that are more biologically plausible than existing methods and accurately infer known cell type transitions.

## 2 Latent Trajectories and Optimal Transport

Given a time series **X** = (**X**^1^,**X**^2^, …,**X**^*N*^) of cells – where each 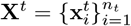 are the cell states for a population of *n*_*t*_ cells – our goal is to learn a collection of trajectories 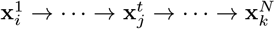 for each cell *i* that link cells from consecutive time points and describe the temporal evolution of individual cells over time. A widely used model in developmental biology to describe these trajectories is Waddington’s landscape [38, 33] which represents each cell state as a point on a landscape with multipotent cells having high developmental potential descending in key furrows of an epigenetic or transcriptomic landscape to a final differentiated population 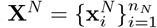 (Fig. 1a).

Importantly, in organismal development cells group into a small number of cell states / cell types. Thus, computing trajectories of individual cells (i.e. a “full-rank” mapping from **X**^*t*^ to **X**^*t*+1^) is likely to overfit the data. Instead, the goal is to derive *latent trajectories* that connect cell types from populations of cells drawn from these cell types without prior knowledge of the cell types. We introduce the problem of discovering latent trajectories from a time series informally in Problem 1 below, which we will later formalize in Problem 2.

### Problem 1

(Latent trajectory inference problem, informal). *Suppose* 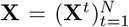 *is a time series with unknown trajectories* 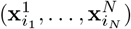 *and latent trajectories* 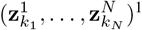 ^1^. *Let* **Q**_*t*_ *be the joint distribution between each* 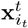 *and its latent state* 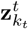, *and assume* 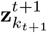 *is generated from* 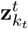 *under transition kernel* 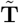.*Given* **X**, *find* **Q** *and* 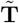.

As one concrete example, sampled single-cell trajectories 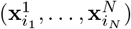 may be well-represented by only a handful of *discrete* latent trajectories 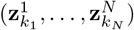,each such latent trajectory indexing a canal in the Waddington landscape. One can sample the initial “cell type” 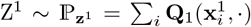 and transition cell types according to transition kernel 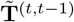 which generates Z^*t*^ conditional on Z^*t*−1^. Given the latent trajectory 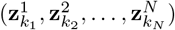,observations are sampled independently at each timepoint: 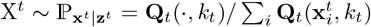. This generates an observed trajectory 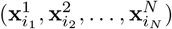 and a latent trajectory 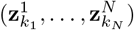.Our focus on single-cell and spatial datasets motivates a formalization of Problem 1 using an empirical distribution over **X** and the framework of optimal transport to identify latent trajectories of *least-action* that are optimal with respect to this principle, which we describe below.

### 2.1 Optimal transport

Optimal transport (OT) is a framework for comparing and aligning datasets, encoded as probability distributions, using distances between the data points to determine the correct alignment. To describe this framework in general, suppose (X,d) is a metric space and let ***X*,3*Y*** ⊂ X be two datasets, 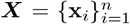 and 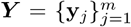. OT represents these datasets as discrete probability measures 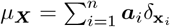 and 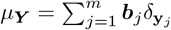, where ***a*** ∈ Δ_*n*_ and ***b*** ∈ Δ_*m*_ are probability vectors (often taken to be uniform), and 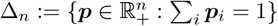 denotes the probability simplex of size *n*. The goal of OT is to find a coupling matrix **P** with *marginals* **P1**_*m*_ = ***a***, and **P**^T^**1**_*m*_ = ***b*** that minimizes a data-driven transport cost. Defining the sets 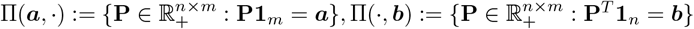,The set of *coupling matrices* or *transport plans* with marginals ***a*** and ***b*** is Π(***a*,*b***) = Π(***a***,·) ∩ Π(·,***b***). Given a cost *c* : X × X → ℝ_+_ and a cost matrix 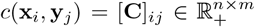,the *Wasserstein problem* is to find **P**^⋆^ ∈ Π(***a*,*b***) minimizing a transport cost over all coupling matrices:

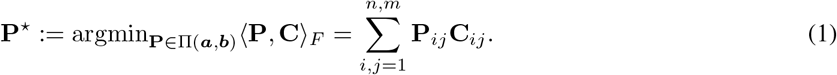

The optimal transport cost itself, ⟨**P**^⋆^,**C**⟩_*F*_ is called the *Wasserstein distance* between ***a*** and ***b***. The Wasserstein problem finds a least-cost alignment between distributions. This has been applied in the seminal Waddington-OT [33] to find a least-cost alignment between single-cell transcriptomics distributions ℙ_*t*_ at different timepoints *t*.

### 2.2 Low-rank optimal transport with LC factorization

Low-rank OT [8, 30, 20, 32, 29] is a way of building interpretable cluster structure into OT by expressing transport plans as a product of low-rank factors performing an implicit clustering. Formally, low-rank OT constrains the *nonnegative rank* rk_+_(**P**) of coupling matrices **P** ∈ Π(***a,b***), defined as the smallest number of nonnegative rank-one matrices summing to **P**. For integer *r* ≥1, let Π_*r*_(***a***,·) := {**P** ∈ Π(***a***,·) : rk_+_(**P**) ≤ *r* },Π_*r*_(·, ***b***) := **P** ∈ Π(·, ***b***) : rk_+_(**P**) ≤ r },and define Π_*r*_(***a*,*b***) := Π_*r*_(***a***,·) ∩Π_*r*_(·, ***b***). The low-rank balanced Wasserstein problem is (1) with Π_*r*_(***a*,*b***) in place of Π(***a*,*b***), and analogously for other marginal constraints and objectives. To solve low-rank OT problems, one must first choose a low-rank factorization of transport plans, with different clustering implications for each choice. Towards the construction of differentiation maps (where the number of cell types changes over time), we choose *latent coupling (LC) factorizations*, first introduced by [20] and recently extended to general costs by [11].

#### Definition 1

(LC factorization). *A* latent coupling (LC) factorization *of a coupling matrix* 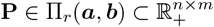 *is a decomposition of the form*

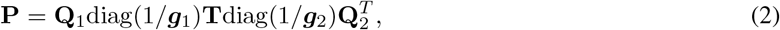

*where* 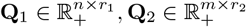 *are the* latent representations *of* ***a*** *and* ***b***, *respectively;* 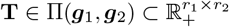 *the* latent coupling; *and* 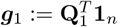 *and* 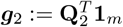 *are the* latent marginals.

One can imagine this decomposing an alignment between two datasets into (1) a map from a distribution over points in the first dataset (**a**) to their cluster distribution (***g***_1_) by **Q**_1_, (2) a matrix of transitions **T** from the first cluster distribution to the second, and (3) a map from the second cluster distribution (***g***_2_) to a distribution over points in the second dataset (**b**) via **Q**_2_ (Fig. 1b). In general, given a dataset 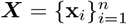 equipped with probability vector ***a*** ∈ Δ_*n*_, a *latent representation* of (***X*,*a***) is a coupling matrix 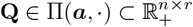,where we typically suppose *r* ≪ *n*. Latent representations **Q** are joint distributions between points **x**_*i*_ and *r* abstract latent points, and can be normalized to be row-stochastic or column-stochastic, with distinct interpretations. The row-stochastic normalization diag(1/***a***)**Q** is a *soft clustering* of the dataset ***X*** using *r* labels, the column-stochastic normalization **Λ** := **Q**diag(1/***g***) is a matrix of emission probabilities giving the likelihood of sampling data points in ***X*** conditional on one of the *r* labels, and column-stochastic 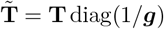 is the cluster-cluster transition kernel.

Let LC_***a,b***_(*r*_1_,*r*_2_) denote the set of LC factorizations (**Q**_1_,**Q**_2_,**T**) with 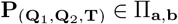 and 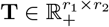,where 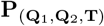 is the product in Definition 1: 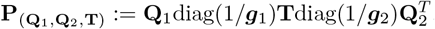. Using LC factorizations, the low-rank balanced Wasserstein problem is

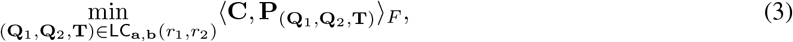

which is equivalent to the objective optimized in [11].

## 3 Hidden Markov Optimal Transport

We extend the inference of latent transitions in the LC-factorization for pairwise optimal transport to *N* > 2 timepoints. This motivates Problem 2 and Problem 3 below, which generalizes the approach to a time-series of any length *N*.

### 3.1 Temporal clustering using HM-OT

Below, we suppose that our time series 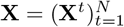 consists of datasets **X**^*t*^ ⊂ X all lying in the same metric space (X,d), and write 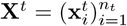.For some cost function *c* : X × X → ℝ_+_, define cost matrices 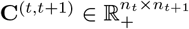 between consecutive time points via 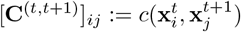.

#### Problem 2

(`HM`-`OT` problem). *Let* 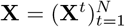 *be a time series of N datasets, where each dataset* **X**^*t*^ *is equipped with probability vector* 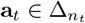.*Given positive integers r*_1_, …,*r*_*N*_, *and cost matrices* 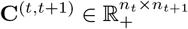,*find a sequence* 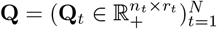 *of latent representations*, and *a sequence* 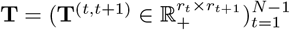 *of latent couplings that minimize:*

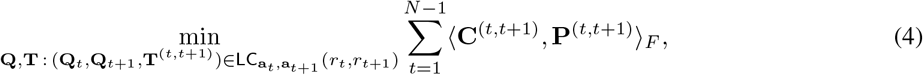

*where* 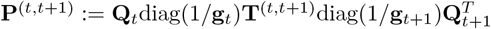.

Above, **P**^(*t,t*+1)^ is defined as a special case of (6), with ***g***_*t*_ the latent marginal of **Q**_*t*_, and **C**^(*t,t*+1)^ are cost matrices between consecutive time points for cost function *c*. Note that the case that *N* = 2, Problem 2 recovers the FRLC objective (3). As with FRLC, it is straightforward to generalize the total distance in the loss, replacing the Wasserstein cost in each term with GW or FGW (to handle spatial transcriptomics data). One can also relax the outer (***a***_*t*_) marginal constraints, as in [10], though we leave a thorough exploration of these relaxed constraints on our output to future work and focus on the balanced case, in which each **P**^(*t,t*+1)^ ∈ Π(***a***_*t*_,***a***_*t*+1_). A solution to Problem 2 is a collection of latent representations 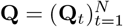 for the time series joined by a sequence of latent couplings 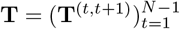;we call this sequence of latent couplings a *differentiation map* (Fig. 1c).

A differentiation map provides the minimum additional input required to describe a joint distribution across **X**, given a corresponding sequence of latent representations **Q**. In particular, we can express the joint distribution of a pair of timepoints *s* < *t* ∈ [*N*] which are not consecutive. First, define latent coupling matrix **T**^(*s,t*)^ by

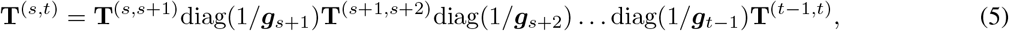

and define the joint distribution of timepoints *s* and *t* from **Q,T** as:

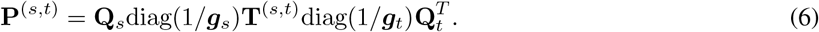

The factorization (5) is (up to the left-most factor) a composition of Markov transition kernels governing latent trajectories between times *s* and *t*. From the hidden-Markov perspective, column-stochastic normalizations **Λ** = **Q**diag(1/***g***) are matrices of emission probabilities. The interpretation of this structured coupling of timepoints **X**^*s*^ and **X**^*t*^ is: first, sample an entry of **T**^(*s,t*)^, a pair of clusters, each at a distinct time point. Then, from each cluster, use the emission probabilities **Λ**_*s*_,**Λ**_*t*_ to sample an element of each timepoint. The emission matrices **Λ**_*t*_ appear naturally in (6), so we will often write 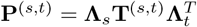 for brevity, with **T**^(*s,t*)^ as in (5). We stress however that the matrices 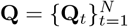 and 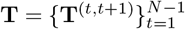 are our variables of optimization in the algorithm presented below.

### 3.2 Computing the optimal Λ-sequence and approximate MAP estimate for HM-OT

We derive an algorithm to learn the set of latent representations and differentiation maps which minimize the objective 4 in `HM`-`OT` problem 2. First, we note that the loss in Problem 3 satisfies an optimal sub-structure property described below, where the optimal loss at time *t* depends only on the optimal loss at *t* − 1. To see this, let ***θ*** = **Λ,T** and define

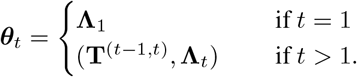

Then the partial sums *ℰ*_*t*_(***θ***_*t*_) express the minimization (4) recursively:

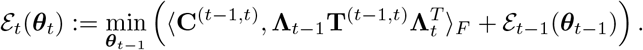

While dynamic programming (DP) principles have been applied to losses over sequences with similar optimal sub-structure properties, applying DP directly to Problem 2 has a number of serious limitations. In the context of HMMs, the Viterbi algorithm is used to infer the maximum-likelihood latent trajectory, under the assumption that the hidden state spaces are finite, and that emission probabilities and transition probabilities are known. In contrast, `HM`-`OT` learns both emission and transition probabilities, which themselves are the sequential variables. Moreover, the variables are real-valued, high-dimensional, and constrained, rendering the discretization necessary to apply a max-sum (Viterbi-type) DP infeasible. We choose instead to approximate the maximum a posteriori (MAP) estimate of **Λ**_*t*_ using a message passing approach. This combines a forward-time “*α*-message” with a reverse-time message “*β*-message” to learn a consensus latent representation 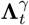 from the full time series.

### 3.3 Forward, backward and consensus passes

The consensus γ-representation is conditional on the past (α) sequence **X**^1^, …,**X**^*t*−1^ *and* the future (β) sequence **X**^t⨥1^, …,**X**^*N*^. In S4.1.2, we derive Algorithm 1 to approximate the MAP estimate 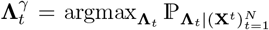 which yields a least-cost clustering at each timepoint *t*, given all other time-points 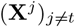.We note that this is distinct from finding the least-cost *sequence* 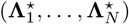 which a dynamic-programming approach would yield.

Like the HMM or Kalman filter, the estimate of the MAP requires the defining a probability density (likelihood) over the variables of optimization. In our case, as we optimize a general energy function ℰ: **Λ**_*i*_ ×**T**^(*i,i*+1)^ →ℝ, this corresponds to a Boltzmann density on our energy

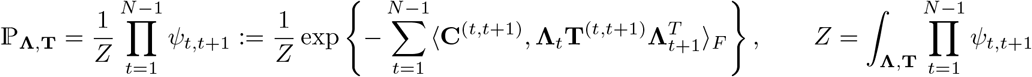

where *ψ*_*t,t*+1_ exponentiates each pairwise cost. We require a recursive set of integrals over future and past couplings to yield a marginal over each **Λ**-triple, 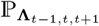.We approximate each integral over the space of couplings by placing a *δ*-measure on the most-likely future and past couplings 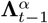 and 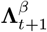,corresponding to the *ground-state* in each direction. This is the well-known *zero-temperature approximation*, where for temperature *τ* one takes the limit

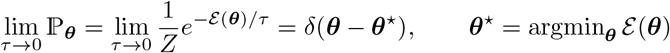

Using this, we can channel information from the past and future sub-sequences with a pair of *α,β*-passes, avoiding an integration over the space of **Λ,3T**. The *α*-pass (Algorithm 2) computes 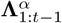 as greedy forward-direction variables and the *β*-pass (Algorithm 3) 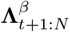 as greedy backward-direction variables. Conditional on a forward message 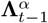 and backward message 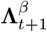,we compute a smoothed variable 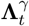 which marginally minimizes the summation objective 2 by minimizing a loss defined on 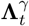 and costs over the local triple of timepoints *t*−1,*t,t*+1. The dependence on the full sequence (**X**^1^, …,**X**^(*t*−2)^), (**X**^(*t*+2)^, …,**X**^*N*^) efficiently factors through 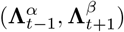 alone, allowing one to tractably compute MAP estimates for massive datasets **X**^(*i*)^ over long sequence lengths *N*. As Algorithm 1 computes smoothed latent representations 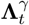 as MAP estimates, the differentiation map is learned in a final completion step. The last step of Algorithm 1 computes optimal transitions between the latent representations to yield differentiation maps between the approximated MAP clusters. By the decoupling of the **T** matrices given **Λ** fixed, this constitutes a simple pairwise OT problem for 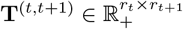 given **Λ**_*t*_,**Λ**_*t*+1_. In fact, this last pass coincides with the fully-supervised variant of `HM`-`OT`, and we refer to (8) for the relevant optimization.

#### Algorithm 1

`HM`-`OT`

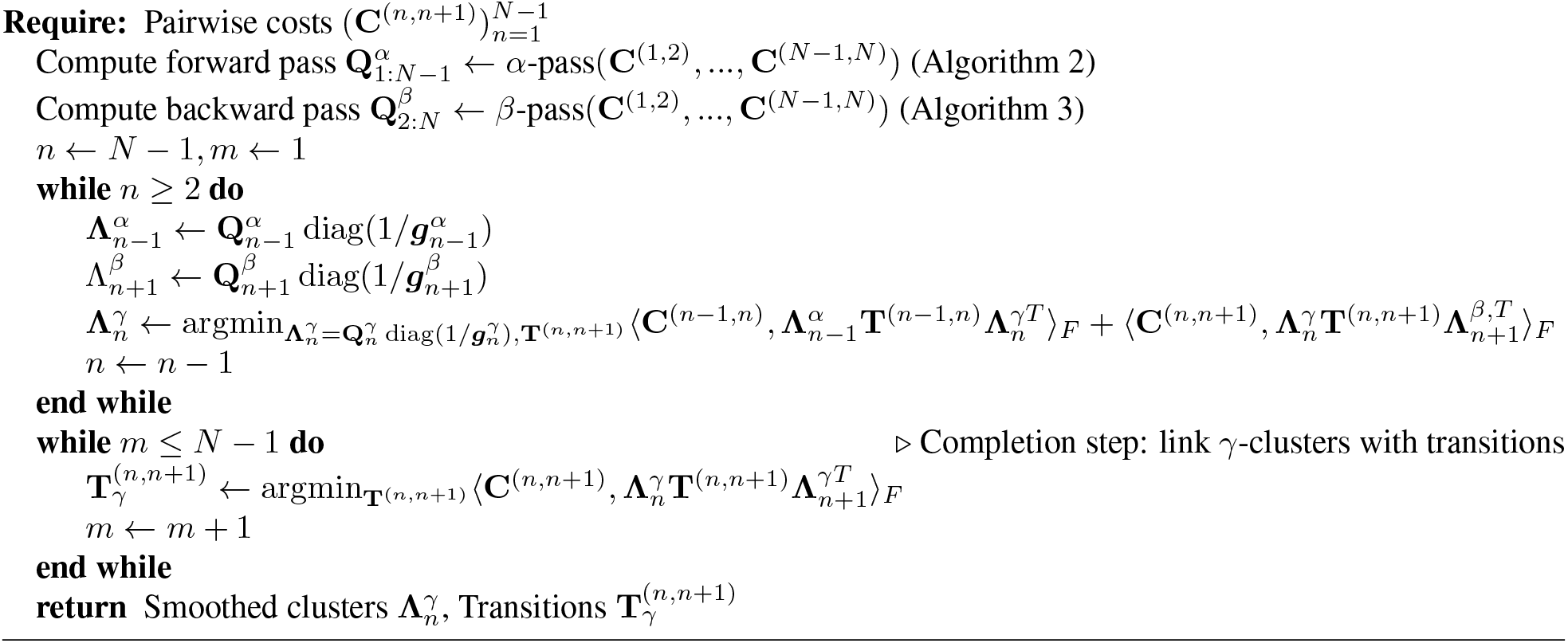

### 3.4 Co-clustering the output of HM-OT

We introduce two approaches to cluster the time series **X** using `HM`-`OT` output. The first, *maximum-likelihood* co-clustering, finds the most likely clusters at each time-point and relates clusters across time-points through the latent coupling matrix. The second, *ancestral* co-clustering, factors the `HM`-`OT` output so that clusters are consistent across time. The former is valuable for identifying cell types and differentiation events. The latter is phylogenetic in nature, allowing one to trace cell types at any intermediate slice to their ancestors and descendants throughout the time series.

#### Definition 2

(Maximum-likelihood clustering). *Given latent representations* 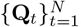,*a maximum-likelihood co-cluster* 𝓁^⋆^(*i*) *for a point i* ∈ [*n*_*t*_] *at time t is defined as the most-likely cluster assignment under* **Q**_*t*_:

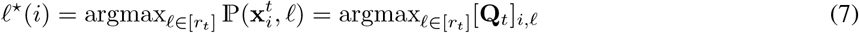

*The clusters at times t,t* + 1 *are put into (soft) correspondence by the cluster-cluster mapping* 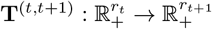.

One can also cluster using a differentiation map by pulling back the mass of each descendant cell to its most likely ancestor, and using the latent cluster of the ancestor as its co-cluster. We call this *ancestral clustering*, defined in Section S5.2 and show one can co-cluster ancestor and descendant cells (Figure S2,S3). Unlike maximum-likelihood co-clustering, which depicts differentiation, ancestral co-clustering depicts “phylogenetic” relationships.

### 3.5 Learning differentiation maps from fixed cell type labels

In some applications, cluster labels for each sample are known; e.g. in a single-cell or spatial dataset, cell types may be annotated using marker genes or a reference dataset of cell types [3, 21, 39], leading to the Supervised `HM`-`OT` problem.

#### Problem 3

(Supervised `HM`-`OT` problem). *The supervised* `HM`*-*`OT` *has an equivalent objective to 2*, *but with any subset* I ⊂ [*N*] *of the latent representations* (**Q**_*i*_) |_*i*∈I_ *fixed, and with the* **T**^(*i,i*+1)^ *and* (**Q**_*i*_) |_*i*∈[*n*]\I_ *remaining free-variables*.

We offer a solution to the Problem 3 and show in S2 how any input clustering at timepoint *t* has a unique latent representation **Q**_*t*_. Thus, fixing cluster assignments for all *t* ∈ [N] uniquely determines all **Q**_*t*_, hence ***g***_*t*_ and **Λ**_*t*_. When all **Q**_*t*_ variables are frozen according to cell type annotations, Problem 3 decouples in time and reduces to a sequence of *pairwise* OT problems for the optimal latent couplings **T**^(*t,t*+1)^, as in the last step of Algorithm 1, reducing to:

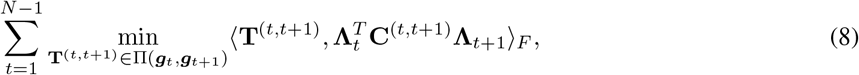

noting the inner products in (8) are equivalent to those of (4). Thus, given annotated cell types as input, one may still find the differentiation map between them using `HM`-`OT`.

### 3.6 Evaluation metrics for differentiation maps

We adopt a widely-used metric to gauge the quality of a differentiation map 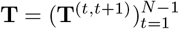.

#### Pointwise Mutual-Information (PMI) for Differentiation Maps

Another measure of the quality of a differentiation map is the deviation between the probability of a transition between cell types a to b and the probability of the same transition under the independence model. Specifically, given the joint distribution **T**^(*s,t*)^, defined in 5, and marginal distributions ***g***_*s*_, ***g***_*t*_ between any pair of timepoints *s, t*, we define the deviation using the *pointwise mutual information*, an information-theoretic metric widely used in machine-learning [15]:

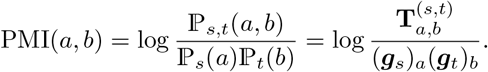

The normalized PMI (NPMI) scales PMI to be in [−1,+1]: where +1 indicates perfect correspondence between a and b, 0 indicates independence, and −1 indicates that a and b for never co-occur.

## 4 Experiments

### 4.1 Synthetic Experiments

We illustrate the advantages of `HM`-`OT` on two synthetic experiments (Fig. S1**a**,**3c**) in Supplement S7.1. First, we demonstrate that `HM`-`OT` more accurately clusters data generated from an external field such as the Waddington gradient (Fig. S1**a**,**3b**) outperforming *k*-means clustering applied independently to each time point (`HM`-`OT` adjusted mutual information AMI = 0.873, *k*-means AMI = 0.393). Second, we demonstrate the distinction between low-rank OT which co-clusters only 2 consecutive timepoints and `HM`-`OT` which simultaneously clusters trajectories with ≥ 2 timepoints (Fig. S1**c**,**3d**). See SupplementS7.1.1,S7.1.2 for details.

### 4.2 Spatial-transcriptomics of Zebrafish development

We evaluate `HM`-`OT`’s ability to derive differntiation maps from spatial transcriptomics data by analyzing a dataset of zebrafish embryogenesis [21] that includes Stereo-Seq spatial transcriptomics data from multiple timepoints of embryonic zebrafish from stages 3.3hpf to 24hpf. We focus our analysis on three time points: 10hpf, 12hpf, and 18hpf. We first run `HM`-`OT` in supervised mode using the cell type annotations from [21], deriving a differentiation map shown in (Fig. 2a). Many of the transitions between cell types in this map do not agree with the known biology of zebrafish development including: notochord at 10hpf transitioning to somite at 12hpf; adaxial at 12hpf transitioning to notochord at 18hpf; and periderm transitioning to segmental plate/tail bud (Fig. 2a). Examining the spatial location of these annotated cell types (Fig. 2b), we observe that the notochord disappears at 12hpf (only 10 isolated spots out of 2081 total spots), but the notochord is a spatially coherent region at 10hpf and 18hpf (Fig. S6). This suggests that errors exist in the cell type labels, which may correspondingly affect the inferred map of cell type differentiation.

**Figure 2:**
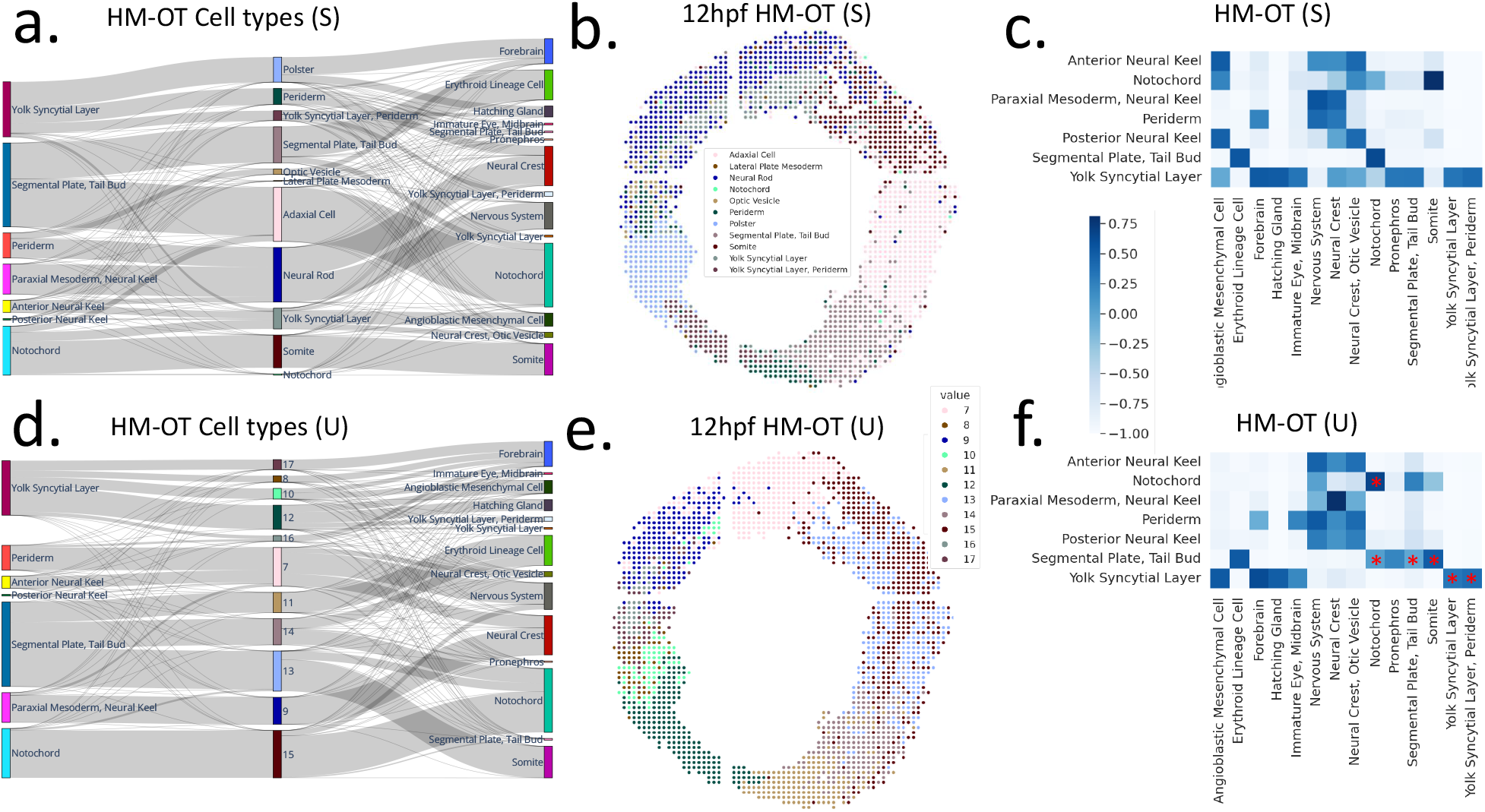
**(a)** Differentiation map output by `HM`-`OT` supervised with annotated 12hpf cell types [21], **(b)** Cell types for the 12hpf timepoint, **(c)** NPMI values between 10hpf and 18hpf for `HM`-`OT` supervised on annotated cell types. **(d)** Differentiation map output by `HM`-`OT` with unsupervised cell types, **(e)** cell types inferred by `HM`-`OT` for 12hpf without supervision, **(f)** NPMI values with `HM`-`OT` cell types. Stars indicate self-transitions and known biological transitions discussed in Section 4.2 .

To further evaluate the differentiation map, we quantify the deviation of the observed trajectories from independence with the normalized pointwise mutual information (NPMI) value for cell types at 10hpf and 18hpf (Fig. 2c). We find that some well known transitions have low NPMI values (indicating that cell types are rarely on the same trajectory) such as notochord to notochord (NPMI = − 0.107), tail-bud to notochord (NPMI = − 0.033), and segmental plate to somite (NPMI =− 0.965).

Because of the inconsistencies in the annotated cell types noted above, we investigate whether the unsupervised mode of `HM`-`OT` can infer better cell types (and consequently, a better differentiation map) on the 12hpf slice. We find that the differentiation map inferred by `HM`-`OT` unsupervised (Fig. 2d), is more biologically plausible than the differentiation map using the [21] annotated cell types. Specifically, `HM`-`OT` suggests notochord transitions to an intermediate cluster (unsupervised cluster 15 in Fig. 2d), which subsequently transitions to notochord, placing it on a single trajectory. In addition, this intermediate cluster 15 is spatially coherent and in the correct biological location (Fig. 2e). The notochord transition has a large NPMI value (NPMI = 0.665) as do several other known biological transitions (Fig. 2f). For example, the transition from tail-bud to notochord, NPMI = 0.610, is also accurate [2]. Notably, `HM`-`OT` suggests the somite cell type arises from its correct progenitor, segmental plate and tail bud [34], with NPMI = 0.497. We offer all NPMI scores (Table S2,S1) and additional discussion of the differentiation maps, including errors common to both, in Supplement S7.2.3.

We further validated `HM`-`OT` unsupervised cluster 15 as the notochord cell type by examining differentially expressed genes. Our identified cluster 15 has *N* = 333 spots and among the top 5 differentially expressed genes (T-test, Benjamini-Hochberg correction) are Chordin, a key notochord marker, (*chrd*, Z = 6.15) and *ripply1* (Z = 10.17) which is known to be found in the notochord [16]. In contrast, the top 5 differentially expressed genes for the notochord cell type at 12hpf using the annotations from [21] are *cirbpb* (Z = 3.32, involved in cold-response), *phf5a* (Z = 2.36, a splice factor), *rps3a* (Z = 2.26, a ribosomal protein), *rpl39* (Z = 2.17, a ribosomal protein), and *prpf4bb* (Z = 2.09, an mRNA processing factor). None of these genes are known notochord markers.

For additional validation, we performed differential expression analysis on `HM`-`OT` clusters that are spatially adjacent to `HM`-`OT` cluster (15), which according to the annotations of [21] is a subset of the adaxial and somite cell type (*N* = 438,265 spots). This region closely matches the region given by `HM`-`OT` clusters (13) and (15). In addition to (15) being a more likely candidate for the notochord cell type, we propose (13) as a better candidate for adaxial. The latter is suggested by the `HM`-`OT` differentiation map, placing (13) on a segmental plate - somite trajectory (Fig. 2d). In Supplement S7.2.2 we show that cell type 13 has muscular proteins characteristic of adaxial and that these proteins are much more weakly expressed in cell type 15. This result recapitulates developmental anatomy – the notochord is a rod-like structure which supports the neural tube and is surrounded by adaxial cells [24, 5]. Cell types 13 and 15 are spatially adjacent, with cell type 15 being more central, and cell type 13 flanking it on each side. Notably, the cell types of [21] relied on the use of a small number of marker genes, while the `HM`-`OT` cell types were inferred from whole-transcriptome expression. This suggests the unsupervised framework of `HM`-`OT` is able to infer cell types without requiring a panel of pre-annotated markers.

### 4.3 HM-OT differentiation map from a time series of whole-mouse embryogenesis

We demonstrate the scalability of `HM`-`OT` by analyzing the mouse organogenesis spatiotemporal transcriptomics atlas (MOSTA) dataset of [3]. This dataset contains spatial gene expression (Stereo-Seq platform) from entire mouse embryo at one day intervals between the time-stages of E9.5 to 16.5 with numbers of spatial locations ranging from 5,913 to 121,767. We compare the differentiation map computed by `HM`-`OT` with the map produced by `moscot`, a method for aligning pairs of spatial transcriptomics slices using optimal transport [17]. To our knowledge `HM`-`OT` and `moscot` are the only current methods that generate differentiation maps from spatiotemporal transcriptomics data and scale to the size of the MOSTA dataset. Both methods rely on low-rank optimal transport, so for both methods we set the rank of the transport matrix to be equal to the number of cell types between each aligned pair.

We find that `HM`-`OT` is both faster than `moscot` and also scales to larger datasets (Fig. S8a, Table S6). `moscot` runtimes (13.95s on *N* = 5,913 spots for E9.5-10.5 to 25.65s on *N* = 18,408 spots for E11.5-12.5) vary with dataset size (Fig. S8a,b), the runtime of `HM`-`OT` grows negligibly with the size of the dataset (8.98s for E9.5-10.5 to 9.31s on *N* = 121,767 spots for E15.5-16.5). Moreover, `moscot` fails to run for alignments beyond the third pair of timepoints and fails to return a transport matrix past the second pair of timepoints (Fig. S8a). While `moscot` uses the low-rank factorization of **P** = **Q** diag(1/**g**)**R**^*T*^, this factorization does not include a latent transition matrix and the latent representations are not returned directly. Thus, one must return **P** in full to derive the differentiation map **T**. For the third pair of timepoints (E11.5 to E12.5) **P** is a matrix with 1.5-billion entries which exceeds GPU memory. In comparison, `HM`-`OT` computes the differentiation map without directly outputting **P**.

We evaluate the differentiation maps returned by `HM`-`OT` (Fig. 3a) and `moscot`(Fig. 3b), generating differentiation maps for both methods supervised by the cell type annotations in [3]. We quantify the difference between the two maps using the NPMI value defined in Section 3.6. We observe that `HM`-`OT` has a wide range of NMPI values with values either near +1 indicating that two cell types lie on a common trajectory or values near -1 indicating they will never share a trajectory (Fig. 3c). In contrast, `moscot` NPMI values are generally close to zero indicating near independence. Across the three timepoints (E9.5, E10.5, and E11.5) where both methods were able to run, `HM`-`OT` captures much stronger transitions between the same cell type than `moscot`. For all 8 cell types which recur from E9.5 to E11.5, `HM`-`OT` differentiation map has higher NPMI values than `moscot`(Fig. 3d). Moreover in the `HM`-`OT` map every cell type is substantially more linked to itself over time compared to `moscot` with a minimum NPMI = 0.176 ≫ 0. As an example, from E9.5 to E11.5 liver has NPMI = 0.713 (`HM`-`OT`) and NPMI = − 0.0475 (`moscot`), branchial arch has NPMI = 0.751 (`HM`-`OT`) and NPMI = 0.144, heart has NPMI = 0.736 (`HM`-`OT`) and NPMI = 0.118 (`moscot`), and brain has NPMI = 0.3507 (`HM`-`OT`) and NPMI = − 0.112 (m`oscot`).

**Figure 3:**
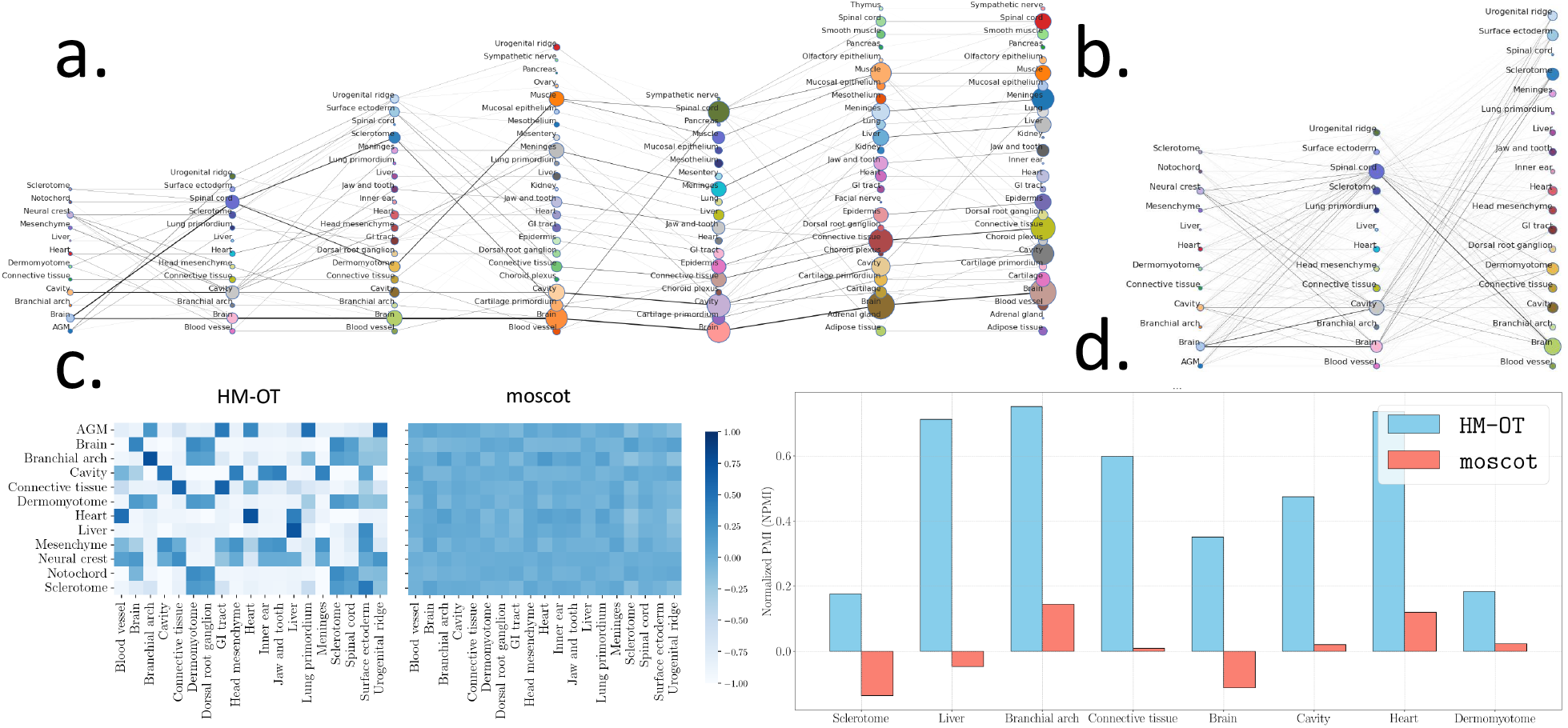
**a**. Differentiation map of mouse embryonic development derived by `HM`-`OT` with respect to the annotations of [3] across timepoints E9.5, E10.5, E11.5, E12.5, E13.5, E14.5, and E15.5. **b**. map of mouse differentiation inferred by `moscot` for E9.5, E10.5, and E11.5 (the map for E12.5 and above exceeded GPU memory). **c**. NPMI scores for all cell types for transitions inferred by `HM`-`OT`and `moscot`. **d**. Comparison on NPMI values for cell types which recur at E9.5 and E11.5 .

Beyond the E9.5-11.5 timepoint, `HM`-`OT` identifies well-known trajectories between differing cell types (Fig. 3a,c). For example, `HM`-`OT` identifies the urogenital ridge transitioning to ovary (NPMI = 0.560, E11.5-12.5) [35], the lung primordium transitioning to lung (NPMI = 0.927, E12.5-13.5) [23, 13], the dermomyotome transitioning to muscle (NPMI = 0.968, E11.5-12.5) [7], sclerotome transitioning to cartilage primordium (NPMI = 0.866, E11.5-12.5) [18], and surface ectoderm transitioning to the epidermis (NPMI = 0.705, E11.5-12.5) [27]. In cases where the cell type annotations are insufficiently resolved to distinguish local ectodermal versus mesodermal regions, one still finds connections between spatially co-localized and functionally related cell types such as the notochord and spinal cord as well as spinal cord and dorsal root ganglion [9]. In summary, `HM`-`OT` recapitulates known developmental trajectories in massive datasets, enabling the automated discovery of differentiation maps at once intractable scales.

Finally, to showcase the advantages of `HM`-`OT`, we visualize growth rates between E9.5-E15.5, derived as described in [10] (Fig. S7), the full map up to E16.5 (Fig. S4), and the co-clustering of mouse-embryonic dataset with both annotated and unsupervised clusters (Fig. S2).

## 5 Discussion

We introduce Hidden-Markov Optimal Transport (`HM`-`OT`) to learn latent temporal trajectories from transcriptomics data by simultaneously identifying latent representations of cells and transition matrices which link these representations across any number of timepoints. `HM`-`OT` can learn biologically plausible cell type differentiation maps, scaling to massive datasets out of reach for other methods. `HM`-`OT` has some limitations which are directions for future work. First, our algorithm computes an approximation of the maximum a posteriori (MAP) sequence of latent representations, and further investigation of scalable approaches to compute the optimal sequence are desirable. Second, our algorithm uses a non-convex objective and depends on hyper-parameters, most notably the number of cell types at each timepoint. Automated selection of these parameters and model selection are important subjects for future work. Finally, while we focused here on applications to spatiotemporal transcriptomics data, `HM-OT` is applicable to longitudinal single-cell transcriptomics data, and future work includes the evaluation of `HM-OT` on these data. We believe `HM`-`OT` and tools that build on it will enable mass-scale unsupervised discovery of the differentiation maps governing development.

## Acknowledgements

This research was supported by NIH/NCI grant U24CA248453 to B.J.R. J.G. is supported by the Schmidt DataX Fund at Princeton University made possible through a major gift from the Schmidt Futures Foundation.

## Supplement

### S1 Optimal transport background

Let (X,d) be a metric space, with vector space structure, and let 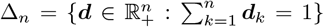 be the set of probability vectors of size *n*. In computational optimal transport (OT), one encodes two datasets ***X*,3*Y*** ⊂ X,

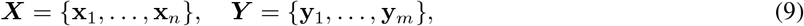

with |***X***| = *n*,|***Y*** | = *m* as discretely supported measures 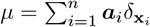 and 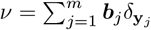 which are represented by probability vectors ***a***∈ Δ_*n*_ and ***b*** ∈Δ_*m*_. A joint density, or alignment, between **a** and **b** is called a *coupling*, which we define below.

#### Definition 3

(Coupling matrix). *The set of* coupling matrices *or* transport plans *between* ***a*** *and* ***b*** *is:*

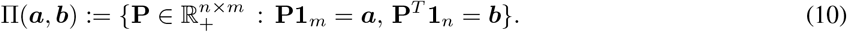

We call ***a*** and ***b*** the *left* and *right marginals* of **P** ∈ Π(***a*,*b***). Using a *cost matrix* **C** derived from the data ***X*,*Y***, one seeks a transport plan **P**^⋆^ optimal with respect to an objective ℋ which is a differentiable function of this alignment.

#### Wasserstein distance (EMD)

Given cost matrix 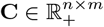 where **C**_*ij*_ = *c*(**x**_*i*_,**y**_*j*_) for *c*(·,·) : X × X → ℝ_+_, the *Wasserstein cost* of transport plan 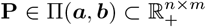 is 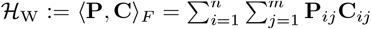. The optimal value of the Wasserstein cost is the *Wasserstein distance between* ***a*** *and* ***b*** *with respect to* **C**.

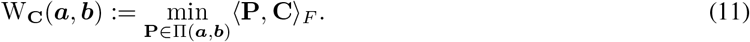

When X is Euclidean space ℝ^d^, and when *c* : X × X → ℝ is the squared-Euclidean distance, (11) defines the squared *2-Wasserstein distance* between ***a*** and ***b***. Wasserstein distance, in particular the 2-Wasserstein distance, is synonymous with *earth-mover’s distance (EMD)*, as the objective value (11) can be interpreted as work or kinetic energy depending on the cost matrix **C**. In typical biological applications, X is taken to be ℝ^*g*^, and where *g* denotes either the number of genes (the dimension of gene expression space) or the number of principal components of the expression vectors that are kept. In such applications, e.g. [33], cost function *c* is usually chosen as the Euclidean distance in this expression space. While we specialize to the Euclidean distance in the present work, we note that all OT problems stated (including the `HM`-`OT` problem statement, Problem 2) extend to general costs, which `HM`-`OT` can handle.

#### Gromov-Wasserstein and other objectives

When datasets ***X*,*Y*** have additional geometric structure (e.g. in spatial transcriptomics), the transport cost (1) can be augmented with a *Gromov-Wasserstein (GW)* term, using intra-dataset distance matrices 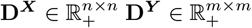 derived from the spatial coordinates of ***X*,*Y***. The *Gromov-Wasserstein* cost of any **P** ∈ Π(***a*,*b***), according to **D**^***X***^, **D**^***Y***^ is:

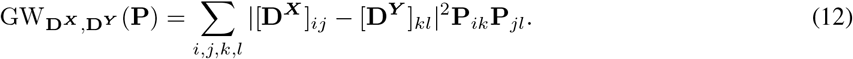

This cost quantifies the metric distortion between ***X*,*Y*** and the optimal 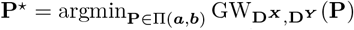 offers a geometric correspondence between ***X*,*Y***. The *fused Gromov-Wasserstein (FGW) problem* finds a transport plan minimizing a convex combination of Wasserstein and Gromov-Wasserstein costs

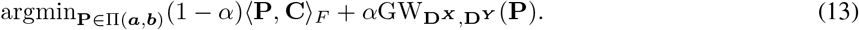

Several works [40, 22, 17, 10] have used the FGW problem to align pairs of spatial transcriptomics datasets in different contexts. Other extensions, such as *semi-relaxed* and *unbalanced* OT, relax constraints on the marginals with a lower semi-continuous *φ*-divergence soft-penalty, such as KL. Such soft-penalties are more outlier-robust and can model growth and death. In particular, [33, 17] use unbalanced OT with curated marker genes to adjust the marginal ***a*** to account for growth and death over time, and [10] learns unsupervised growth rates using semi-relaxed OT.

It is possible that the two datasets ***X*** ⊂ X,| ***X***| = *n* and ***Y*** ⊂ | 𝒴|, = *m* live on distinct spaces, X ≠ Y. Though we assume X and Y are each equipped with a metric, respectively d_X_ and d_Y_, there may not be a common metric d with which to compare *x*_*i*_ ∈ 𝒳 across the datasets to some *y*_*j*_ ∈ 𝒴. In this setting, we no longer have 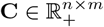.However, one can still form *intra-dataset distance matrices* 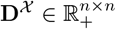 and 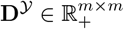 from the data (9) via

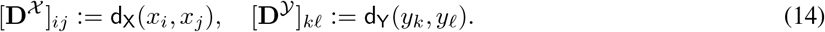

These are used to define the *Gromov-Wasserstein objective*

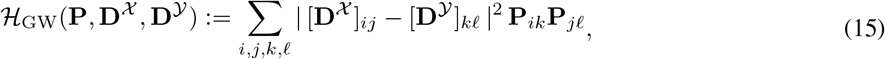

and the *Gromov-Wasserstein problem* for ***a*** ∈ Δ_*n*_, ***b*** ∈ Δ_*m*_ is the minimization of (15) over **P** with ℋ = ℋ_GW_(**P,3D**^𝒳^, **D**^𝒴^). The minimum of the objective (15)

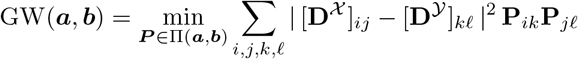

quantifies the minimum metric distortion incurred over all samples from **P**, where each sample from **P** is an (*x,y*)-pair in 𝒳 ×𝒴

In other instances, one has access to both d and d_X_,d_Y_. In the simplest case, all three metrics coincide. However, it is possible that data 𝒳,𝒴 are additionally paired with accompanying features from different modalities, possibly distinct across the two datasets. In this case, with more structured data (and also in the simplest case), d_X_ and d_Y_ can be used to define intra-dataset distance matrices **D**^𝒳^, **D**^𝒴^ for these modalities, as above in (14), and one can combine the Wasserstein and Gromov-Wasserstein objectives through a hyperparameter *α* ∈ (0,1) [36]:

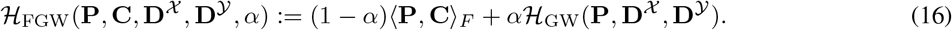

Objective (16) is called the *fused Gromov-Wasserstein (FGW) objective*. FGW objectives have been used to align spatial transcriptomics data [40, 22, 17], and variants of FGW have been introduced [10] that include a sum over triples of points, effectively upweighting a subset of the terms in the sum (15).

### S2 Clustering and co-clustering

#### Background on clustering

Let (X,d) be a metric space with vector space structure, and let 𝒳 = {*x*_1_,…, *x*_*n*_} ⊂ X be finite a set. Suppose we are given a dissimilarity function *d* : X × X → ℝ _+_ which satisfies reflexivity so that *d*(*x,x*) = 0, and symmetry so that *d*(*x,y*) = *d*(*y,x*) ∀*x,y* ∈ X. Define [**C**]_*ij*_ := *d*(*x*_*i*_,*x*_*j*_) to be this dissimilarity function evaluated between all pairs of points in the finite set 𝒳.

##### Definition 4

(Clustering). *Define P*(𝒳) *to be the set of all partitions of a set* 𝒳= {*x*_1_,…,*x*_*n*_}. *A* clustering function on 𝒳 ⊂ X *is a map F* : **C** → *P*(𝒳). *A* clustering *is the corresponding partition F*(**C**) = *π* ∈ *P*(𝒳) *of the dataset*.

Generalizations of clustering often weight the points of 𝒳 non-uniformly, corresponding to a weight vector w placed on the data points. In the context of 𝒳 optimal transport, we specialize to weights on the probability simplex 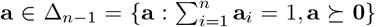,and represent our dataset 𝒳 as a measure supported on X weighted by a, 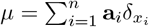. For each clustering function *F*, one can define an additional probability vector g, where 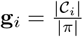 for all clusters in the partition 𝒞_*i*_ ∈ *π*. That is, the density g corresponds to (generally non-uniform) cluster proportions. Trivially, one can form a coupling γ ∈ Π(**a,3g**) by defining an empirical joint measure 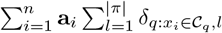.

In a similar sense, we note that any map *F* : **C** → *P*(𝒳) can be equivalently represented by a matrix **Q** ∈ {0,1}^n×|*π*|^ where

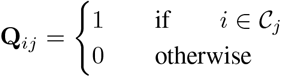

##### Definition 5

(Soft clustering). *A* soft clustering *is a relaxation of 6 where* **Q**_*ij*_ ∈ ℝ _+_ *and* ∑_*j*_ **Q**_*ij*_ = 1.

As such, the definition of soft-clustering in 5 allows fuzzy assignments of points *i* to clusters *j* where the assignment may be split, e.g. between two clusters *j,j*^′^ which are both close to *i*.

##### Definition 6

(Co-clustering). *Define P*(𝒳 ∪ 𝒴) *to be the set of all partitions of the union of a set* 𝒳 = {*x*_1_,…,*x*_*n*_} *and a set* 𝒴 = {*x*_1_,…,*x*_*m*_}. *Let* [**C**_*XY*_]_*ij*_ = *d*(*x*_*i*_,*y*_*j*_) *be a matrix with rows representing* 𝒳 *and columns representing* 𝒴. *A* co-clustering function on 𝒳 × 𝒴 ⊂ X × Y *is a map F* : **C**_*XY*_ → *P*(𝒳 ∪ 𝒴). *A* co-clustering *is the corresponding partition F*(**C**_*XY*_) = *π*_*XY*_ ∈ *P*(𝒳 ∪ 𝒴) *of the row and column clusters of the matrix*.

As before, a co-cluster map *F* can be represented in a matrix form – now with a pair of matrices **Q** ∈ {0,1}^*n*×|*π*|^ and **R** ∈ {0,1}^*m*×|*π*|^ where

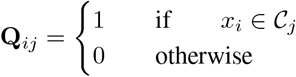

and

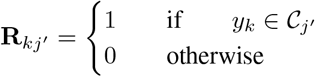

where *x*_*i*_ and *y*_*k*_ are said to be *co-clustered* if *j* = *j*^′^. In the case of co-clustering, one can observe that the cluster proportions allocated to 𝒳 and 𝒴 may not be equivalent. In particular |{*x* ∈ 𝒳 : *x* ∈ 𝒸_*j*_}| ≠ |{*t* ∈ 𝒴:∈ *y* 𝒸_*j*_}|, and *n* ≠ *m* in general. Even more broadly, it may be so that the set of clusters 𝒞_*j*_ representing 𝒳 and 𝒴 are jointly not as reasonable as a pair of sets of clusters 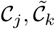 allowed to have differing cluster-proportions allocated to 𝒳,𝒴 with a *joint-density* between the clusters themselves.

##### Definition 7

(Joint-factored Co-clustering). *Define P*(𝒳),*P*(𝒴) *to be the set of all partitions of* 𝒳 = {*x*_1_,…,*x*_*n*_} *and* 𝒴 = {*y*_1_,…,*y*_*m*_}. *Let* [**C**_*XY*_]_*ij*_ = *d*(*x*_*i*_,*y*_*j*_) *be a matrix with rows representing* 𝒳 *and columns representing* 𝒴. *A* joint co-clustering function on 𝒳 × 𝒴 ⊂ X × Y *is a map F* : **C**_*XY*_ → *P*(𝒳),*P*(𝒴),*µ*_*XY*_ *where µ*_*XY*_ *is a joint-density defined on the product-space of partitions P*(𝒳) × *P*(𝒴). Marginal clusterings *correspond to the partitions of* 𝒳 *and* 𝒴, *π*_*X*_ *and π*_*Y*_, *and the* co-clustering *corresponds to partitions in P*(𝒳 ∪ 𝒴) *maximally-likely under the joint law*.

Under this definition, one allows both distinct clusters on 𝒳 and 𝒴, as well as distinct cluster densities 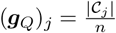 and 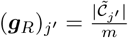 where the difference between clusters is mediated by the joint density, or coupling, in the set

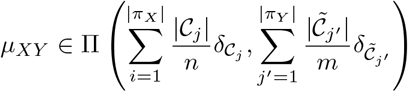

with matrix-realization 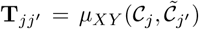 quantifying the joint-density between clusters *j,j*^′^ of datasets 𝒳,𝒴. Having defined joint-factored co-clustering 7 under the assumption of hard 0-1 assignments, we note that there is a direct extension to soft co-clustering, analogous to 5, where the integral constraint is relaxed and one only requires that 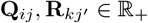 and 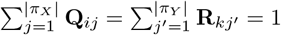.

#### Connection to the LC-factorization

One may encode any input clustering as a unique **Q**_*i*_ matrix of an LC-factorization, and given these **Q**_*i*_ matrices, can solve a sequence of *pairwise* OT problems for the differentiation map alone. Suppose one is given the cluster assignment of each sample *i* to a cell type l for times *t* = 1, ‥,*N* as a map *F* : [*n*_*t*_] → [r_t_] for *r*_*t*_ fixed cell types. Representing the encoding of cells by their type one can form the empirical joint measure, or coupling, given by 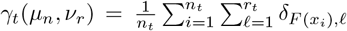 which has the matrix representation of 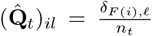. The row-sum represents the proportion of cluster 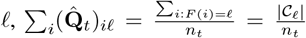 where 𝒞_𝓁_ denotes the set of points assigned to cluster 𝓁. The column sum represents the proportion of spot *i*, taken to be 1/*n*_*t*_ for the empirical coupling, 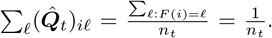. Thus 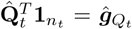 has the cluster-distribution as its entries and 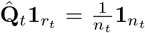 is uniform over points. Thus, having a cluster assignment uniquely determines an encoding of a sub-coupling matrix variable 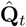.Thus, given marginal clusterings *π*_*X*_,*π*_*Y*_ in 7, one uniquely determines two latent representations for **Q**_*X*_,**Q**_*Y*_, and the matrix realization of a joint-density in 7, 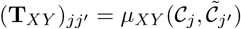, determines a latent coupling as in 1. Thus, a joint-factored co-clustering 7 uniquely determines an LC-parametrization (**Q**_*X*_,**Q**_*Y*_, **T**_*XY*_).

### S3 Temporal clustering

Temporal clustering has been explored in a number of works, such as Dey et al. [6]. We offer definitions of temporal clustering as it pertains to this work and offer a “generative” process by which one can sample trajectories from one’s unlabeled dataset – e.g. having solved for (**Q,T**) using `HM`-`OT` these may correspond to low-cost trajectories with respect to transcriptional distance.

#### Definition 8

(Latent trajectory clustering). *Suppose* (X,d) *is a metric space, and let* **X** = (**X**^1^,…, **X**^*N*^) *be a sequence of N finite subsets of* X, *where* |**X**^*t*^ |= *n*_*t*_. *Let* **x** = (**x**^1^,…, **x**^*N*^) ∈ **X** be any *fixed trajectory, and let Suppose* 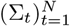 *be finite alphabets with* |Σ_*i*_| = *r*_*i*_. *A* trajectory clustering *is an assignment of any observed trajectory* **x** *in a metric space* X *to a latent sequence*

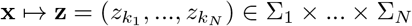

*The discrete sequence* **z** *is the* latent trajectory *of* **x**.

Under this definition, there are 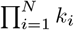 possible latent-trajectory assignments to an observed sequence **x**. In many cases, the set of possible trajectories is much smaller than this – in the case of Waddington’s canalization, Σ_*i*_ represents an alphabet of cell-states on a trajectory where single-cell trajectories may be regarded as sparse in the space Σ_1_ × … × Σ_*N*_. A reasonable assumption to make is that **x**^*i*^ is independent of all **x**^*j*≠*i*^,**z**^*j*≠*I*^ given 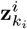, akin to the assumption that on this trajectory **z**, the marginal distribution of **x**^*i*^ depends only on a latent state at time *i*. Under this assumption, one has the following factorization of the joint-density

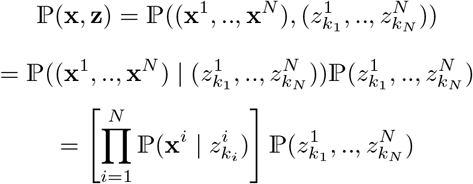

In the case that the trajectories **z** exhibit undirected structure, one may factor the joint over **z** by clique where the joint of the trajectory 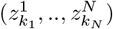 decomposes into a joint over pairs

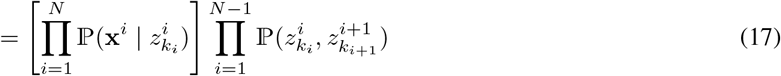

With directed structure over 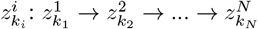, the joint over the trajectory **z** factorizes as a Markov chain

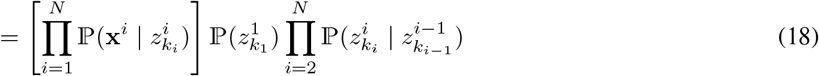

In other words, under our model parameters **Q,T**, 17 the joint on (**x,3z**) has realization given as

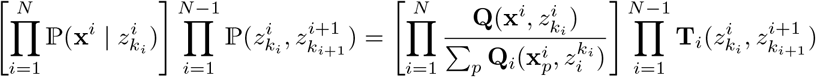

and under 18 the joint on (**x,3z**) has realization

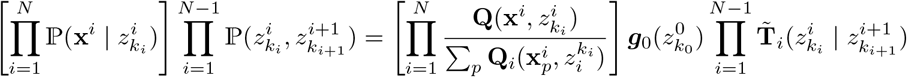

for transition kernel 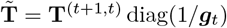.Under this process, one can “generate” latent trajectories within one’s dataset as follows:

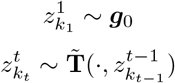

and from the sequence 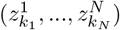,sample a trajectory of points for all *t* ∈ [*N*] as

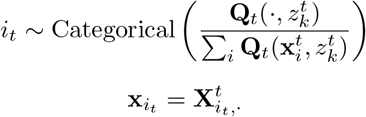

where 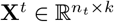 is a data matrix with *n*_*t*_ points (e.g. single-cell transcript vectors) of dimension *k*. Moreover, the question of finding a sparse set of trajectories generating **X** is one of interest, especially in the context of single-cell transcriptomics.

#### Problem 4

(Latent trajectory subset problem, informal). *Suppose* 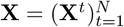 *is a time series where* 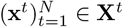 *are associated to latent trajectory*

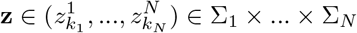

*Identify the subset of latent trajectories* Ω ⊂ Σ_*N*_ *such that* **x** ∈ **X** ⟹ **x** ⟼ **z** ∈ Ω. *Analogously, given a measure µ on* **X** *and a trajectory assignment* 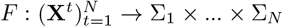,*one seeks a subset* Ω ⊂ Σ_1_ × … × Σ_*N*_ *such that µ* ({**x** : *F*(**x**) ∈ Σ_1_ × … × Σ_*N*_ \ Ω}) = 0 *and µ*(*F*^−1^ (**z**) : **z** ∈ Ω) > 0 *for F* ^−1^(**z**) = {**x** ∈ **X** : *F*(**x**) = **z**}.

For example, Problem 4 seeks a subset of latent trajectories, corresponding to a sequence of latent cell states, which describes the sparse canalization of Waddington’s landscape.

### S4 Solving the Multi-marginal transportation problem

Let **X** = (**X**^1^,…, **X**^*N*^) be a time series of length *N*, with 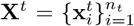,and each **X**^*t*^ equipped with probability vector 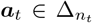.Given couplings **P**^(*t*−1,*t*)^ ∈ Π(***a***_*t*−1_,***a***_*t*_) and **P**^(*t,t*+1)^ ∈ Π(***a***_*t*_,***a***_*t*+1_), let us express their total transport cost across three consecutive timepoints **X**^*t*−1^, **X**^*t*^, **X**^*t*+1^ in terms of random variables (*X*^*t*−1^,*X*^*t*^) ∼ **P**^(*t*−1,*t*)^ and 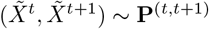:

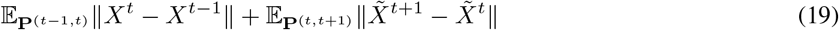

When **P**^(*t*−1,*t*)^ and **P**^(*t,t*+1)^ have no rank constraints, the problem of minimizing (19) over **P**^(*t*−1,*t*)^ and **P**^(*t,t*+1)^ decouples into a sum of independent optimizations. However, we constrain the transport to pass through a *consistent* set of clusters at each timepoint: not only are **P**^(*t*−1,*t*)^ and **P**^(*t,t*+1)^ are low-rank, but we further require their LC factorizations to share a common factor **Λ**_*t*_. To see this, suppose the sets 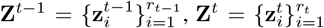,and 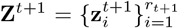 label the clusters at the three time points, and define the following *emission probabilities* from matrices **Λ**_*t*−1_, **Λ**_*t*_, and **Λ**_*t*+1_:

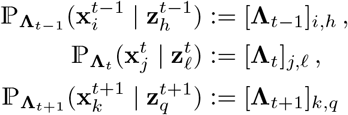

and from latent coupling matrices **T**^(*t*−1,*t*)^ and **T**^(*t,t*+1)^, define the joint distributions over consecutive pairs of cluster labels:

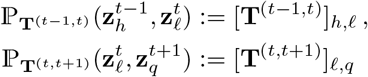

from which, (19) can be expressed as follows:

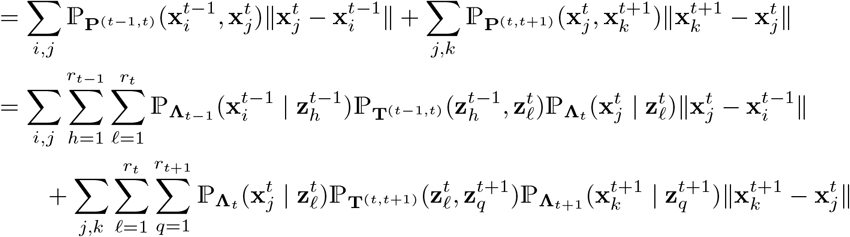

We can further simplify by pulling the common factors of 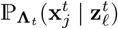.

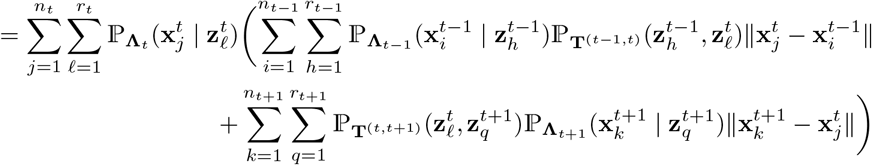

Thus, under our constraints on **P**^1^ and **P**^(2)^, objective (19) reduces to

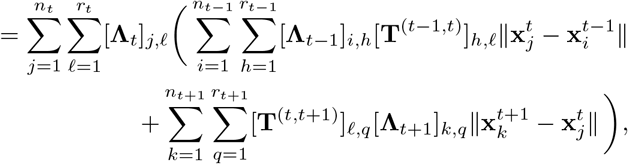

which can be expressed concisely as:

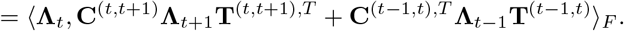

Thus, subject to this low-rank decomposition, to find a clustering which minimizes the total Wasserstein distance traveled across timepoints, the inner clustering must minimize an inner product dependent on its left and right neighbors in the total sequence.

#### S4.1 Optimal substructure and discretized approximations for the optimal Λ-sequence

Let **V** = **Λ,T** denote all optimization variables. Let us group these by timepoint, defining

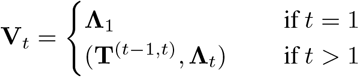

One can express our full loss recursively:

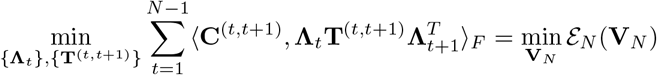

where 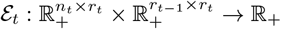 is defined for *t* = 2, …, *N* via:

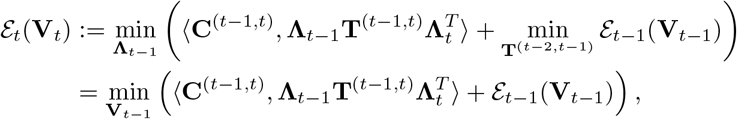

while in the “base” case of *t* = 1, define

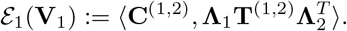

This satisfies the optimal sub-structure property of problems typically solved using dynamic programming. In fact, maximizing a sum over a graph structure is solved by the max-sum DP, of which a sub-case of the algorithm specialized to HMMs is commonly known as the Viterbi algorithm. A number of difficulties make an immediate solution of this problem using standard tabular dynamic programming intractable. The matrices 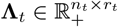 and 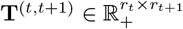 are real-valued, positive matrices. Thus, solving this as a dynamic programming problem requires a discretization of the space which would suffer from the curse of dimensionality, incurring error exponential in the dimension. Moreover, this optimization is heavily constrained: ∀*t* ∈ [1, ‥,*T*] the matrices (**Λ**_*t*_,**Λ**_*t*+1_,**T**^(*t,t*+1)^) must satisfy

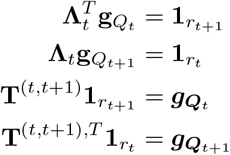

while optimized over all 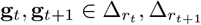.Thus, the discretization of the space would be highly non-trivial. While the problem does satisfy optimal sub-structure by virtue of being a loss over a sequence, being both continuous and highly constrained makes the application of DP infeasible, necessitating approximate or greedy solutions.

#### S4.1.1 Background: decomposing the marginal into forward and backward densities

Our sequence of datasets has the structure of a graph *G* = (*V,E*), with vertices given by the marginal clusterings *v*_*t*_ = {**Λ**_*t*_ } ∈ *V* and edges (*t,t* + 1) ∈ *E* between timepoints in sequence represented by the joint distribution 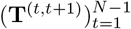.For such graphs with sequential structure, message-passing is a common means of inferring variables which minimize a loss over the graph. We provide relevant background for this, following the presentation in [1]. Suppose one has an undirected chain of nodes (**z**_1_,…,**z**_*N*_). In such a chain one can decomposes the joint probability of ℙ (**z**_1_,…,**z**_*N*_) into cliques consist of pairs of points 𝒞_*i,i*+1_ = {**z**_*i*_,**z**_*i*+1_}, such that the joint probability decomposes into a product of clique potentials *ψ* : 𝒞_*i,i*+1_ → ℝ normalized by a partition function *Z*:

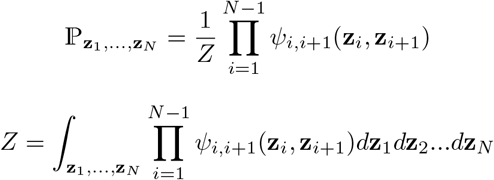

In message-passing, one assumes that the potentials ψ_*i,i*+1_ are given. Using these, message-passing allows for efficient marginalization over all states to yield a distribution on any given node **z**_*n*_ or local connected sequence of nodes **z**_*n*_,…,**z**_*n*+*q*_. It is straightforward to observe that this marginalization is simple to perform from both ends by memoization of “messages” which are passed from both directions. In particular, one may observe

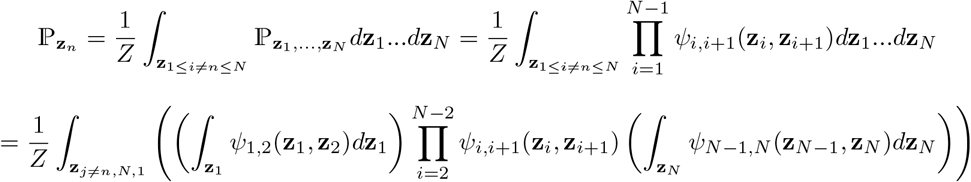

observing that we can compute these integrals and store them as functions of **z**_2_ and **z**_*N*−1_, we renotate the forward and backward messages to be

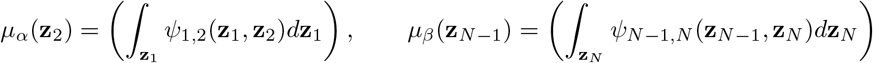

so

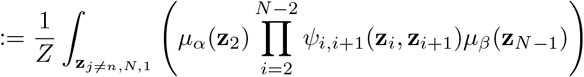

and continue inductively up to node *n*

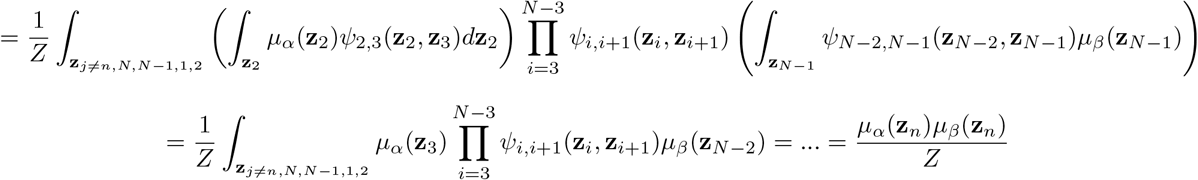

Thus, the marginal on any node 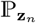 in the graph can be computed using a forward message *μ*_*α*_ and backward message *μ*_*β*_.

#### S4.1.2 Approximating the MAP estimate over triples

##### Proposition 1.

*Supposing* `HM`*-*`OT` *energy function 2 has density given by the Boltzmann distribution*

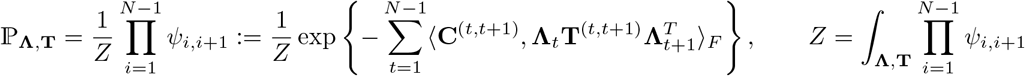

*For pairwise clique energies*, 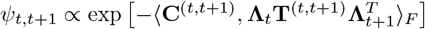, *the MAP estimate over triples* 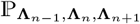 *can be approximated by Algorithm 1 assuming a zero-temperature approximation on the density over* ψ_*n,n*+1_ *in the recurrent integral*

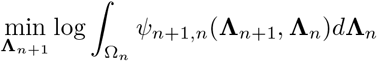

*Proof*. Using the derivation of S4.1.1, one sees that the joint is expressed as

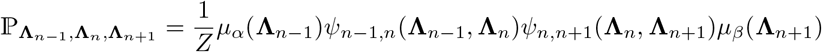

and

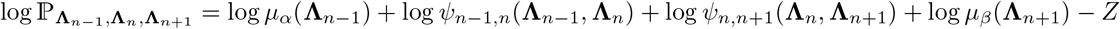

The partition function, which normalizes the Boltzmann exponent, is

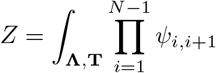

and is constant with respect to **Λ** and **T**. Thus, this reduces in proportionality to

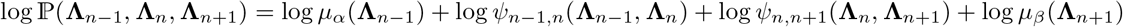

where, for simplicity, the dependence on the transition is left implicit

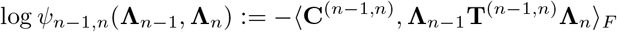

the first core approximation made is as follows

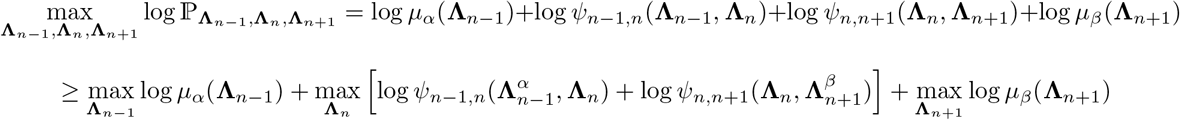

where we maximize a lower-bound on the log-likelihood rather than the log-likelihood itself for

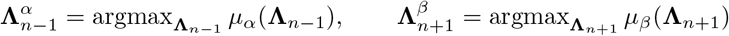

This involves first recursively maximizing a forward message *μ*_*α*_(**Λ**_*n*−1_) and a backward message *μ*_*β*_(**Λ**_*n*+1_), fixing 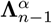 and 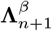 as the associated optimal arguments and then minimizing the **Λ**_*n*_ dependent term as

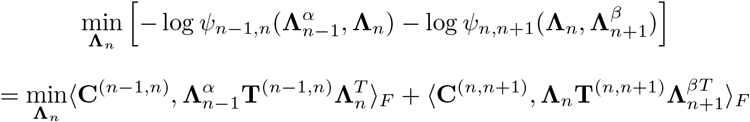

To compute the recursive value for *μ*_*α*_ requires an integration over a space of positive, stochastic matrices **Λ**_*n*_ ∈ Ω_*n*_ subject to the structural constraint that

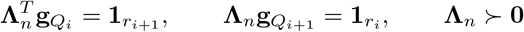

over all 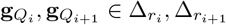.The associated integral is then

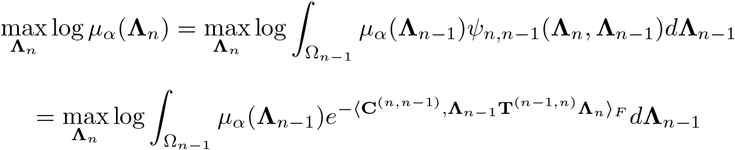

To simplify this, consider the boundary case of *n* = 1 (resp. *n* = *N*):

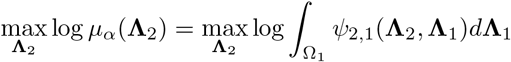

We use the zero-temperature approximation where a Boltzmann distribution is approximated by a *δ*-function on its ground-state

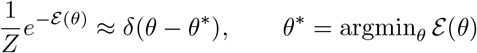

so that annealing the Boltzmann distribution ψ_*n,n*−1_ with a temperature parameter *τ*, by taking *τ* → 0 we approximate the density with a point mass on the most likely configuration (ground-state)

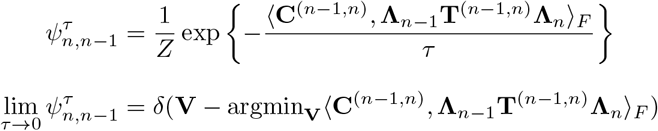

This replaces the intractable to integrate space of Ω_1_ with an integral on a Dirac delta-measure on the ground-state so that

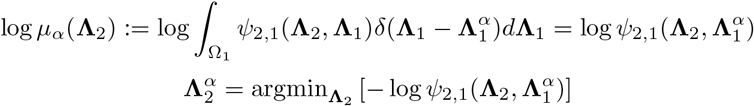

This approximation allows us to compute the forward message *μ*_*α*_ and with identical reasoning the backward message *μ*_*β*_. The recursion as expressed continues this approximation as

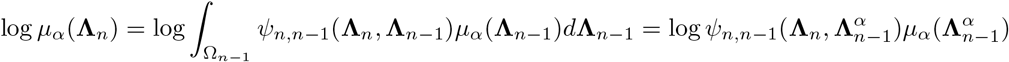

So that

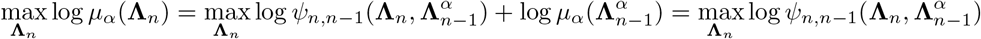

Identical reasoning applies for the backward pass quantities 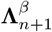 which may similarly be recursively computed starting from time *N*:

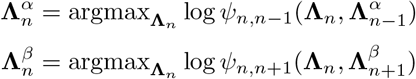

Analogously, if one were to imagine a similar argument on the latent points *z*^*t*^, which are not explicitly variables in our formulation, one would find on our likelihood 17 that

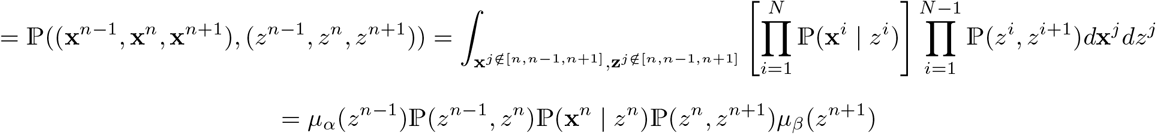

So that the maximization is expressed

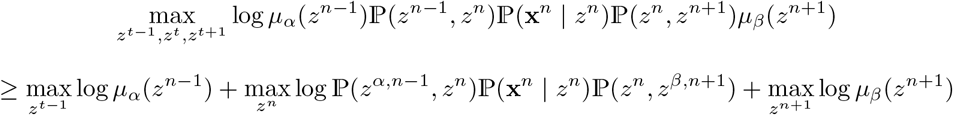

So that given *z*^*α,n*−1^ and *z*^*β,n*+1^ as forward-backward variables one optimizes 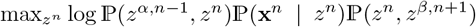 alone, implying that ℙ (*z*^*α,n*−1^,*z*^*n*^) := **T**^(*n*−1,*n*)^, ℙ (*z*^*n*^,*z*^*β,n*+1^) := **T**^(*n,n*+1)^, and ℙ (**x**^*n*^ | *z*^*n*^) := **Λ**_*n*_ are optimized given 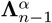 and 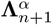 fixed. This similarly implies the step:

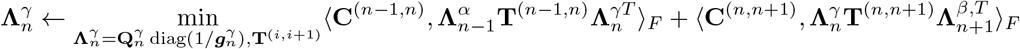

Although a MAP estimate for **Λ**_*n*_ is distinct from the solution in the most likely total sequence 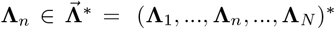,one can still find an optimal set of transition matrices 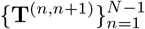 to link them. In particular, given the smoothed matrices 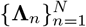, one may connect them in a transport chain directly by performing a final optimization for the optimal transition between each pair of smoothed clusters.

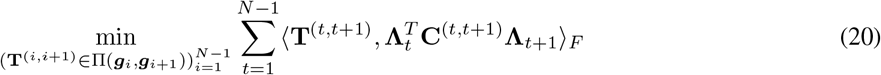

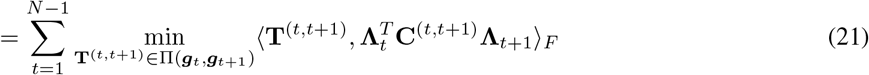

Thus, by the decoupling of the **T** matrices given **Λ** fixed, this constitutes a simple pairwise OT transport problem for 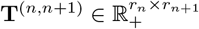 given **Λ**_*n*_, **Λ**_*n*+1_ fixed:

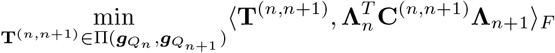

##### Algorithm 2

α-Pass

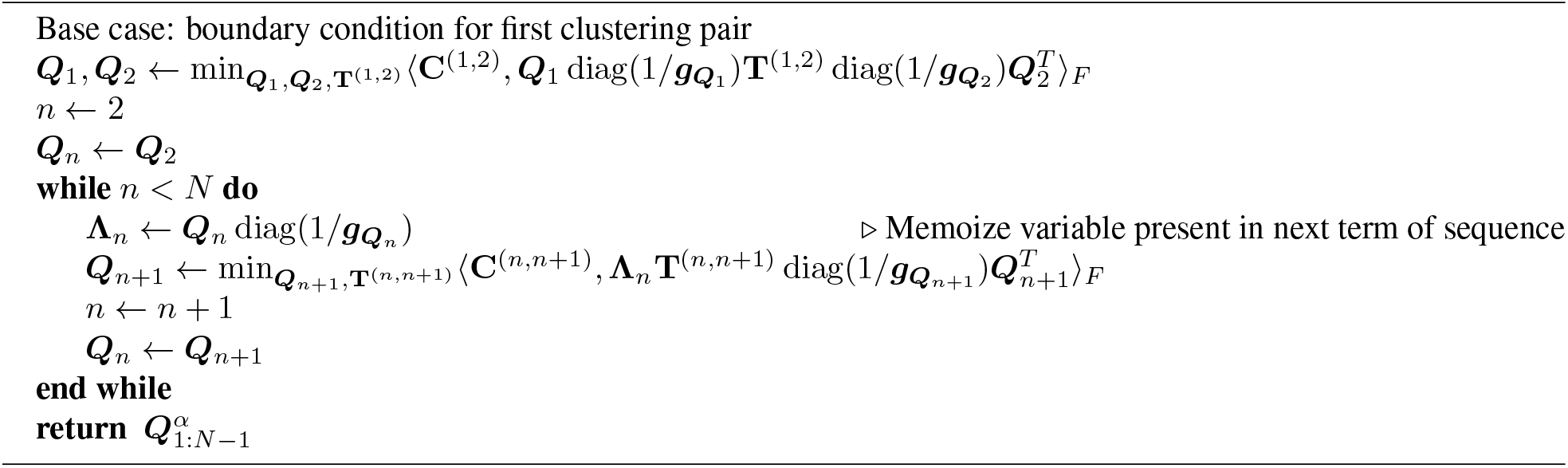

### S4.2 Low-rank factorization of products of cost matrices

The only input to Algorithm 1 are the pairwise distance matrices 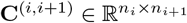 which capture the dissimilarity between a pair of datasets **X**^(*i*)^ and **X**^(*i*+1)^. However, when *n*_*i*_ × *n*_*i*+1_ is a prohibitively large value, it becomes necessary to take a low-rank factorization of the distance matrix itself. Algorithms to do so have been investigated in [14], and applied to OT cost matrices in [30, 31, 32]. In applications involving multi-modal data, such as spatial-transcriptomics, it is often effective to geometrically average the cost matrices 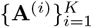 as 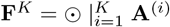 [10]. In S4.2, following previous works which offer the factorization when one takes the Hadamard product of a low-rank matrix with itself [32], we prove Proposition 2 to generalize this to the case one has two distinct low-rank matrices 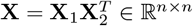 and 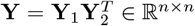 of potentially distinct ranks *d*_1_,*d*_2_ for a factorization of **X** ⊙ **Y**.

#### Algorithm 3

β-Pass

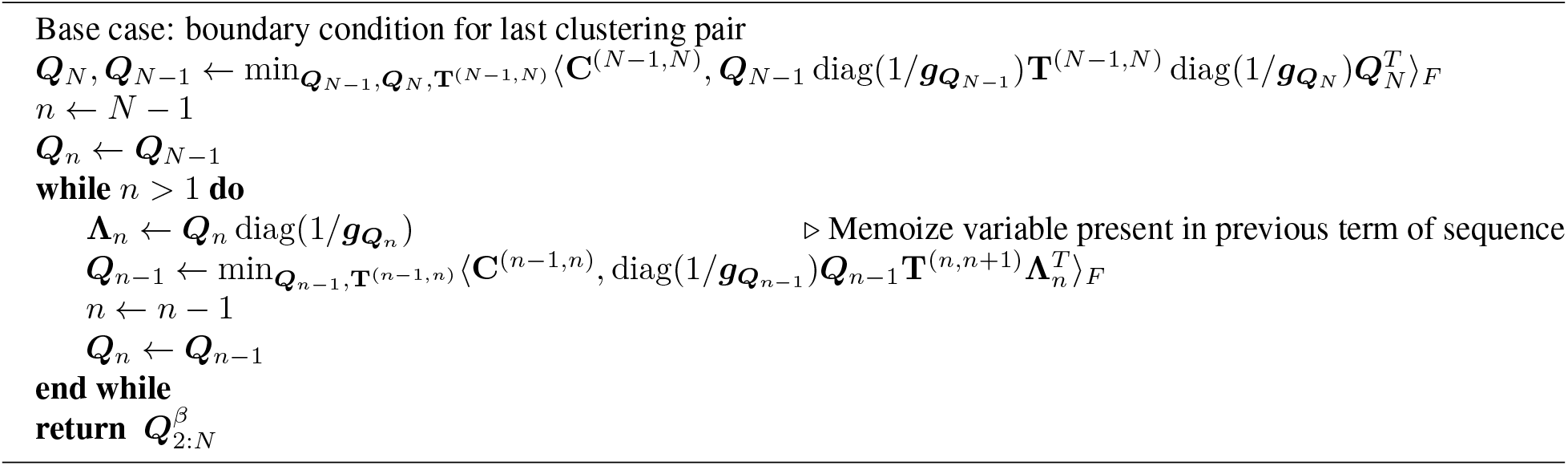

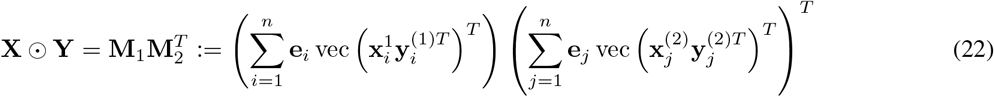

Where 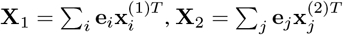, and likewise **Y**_1_,**Y**_2_. Thus, taking 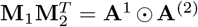 as a base case and continuing inductively, one can geometrically average factored distance matrices 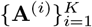 as 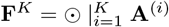

#### Proposition 2.

*Suppose we want to compute* **X** ⊙ **Y** *where* **X, Y** ∈ ℝ ^n×n^ *admit low-rank decompositions of the form* 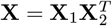 *and* 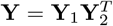 *where* 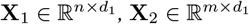 *and* 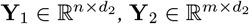.*Suppose we denote:*

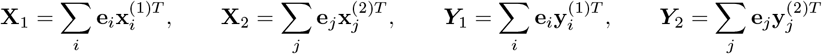

*Then one may use the low-rank factorizations of* **X** *and* **Y** *to factorize their Hadamard product into matrices of dimension* 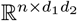

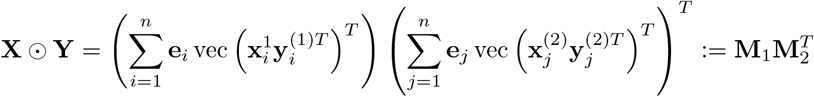

*Proof*.

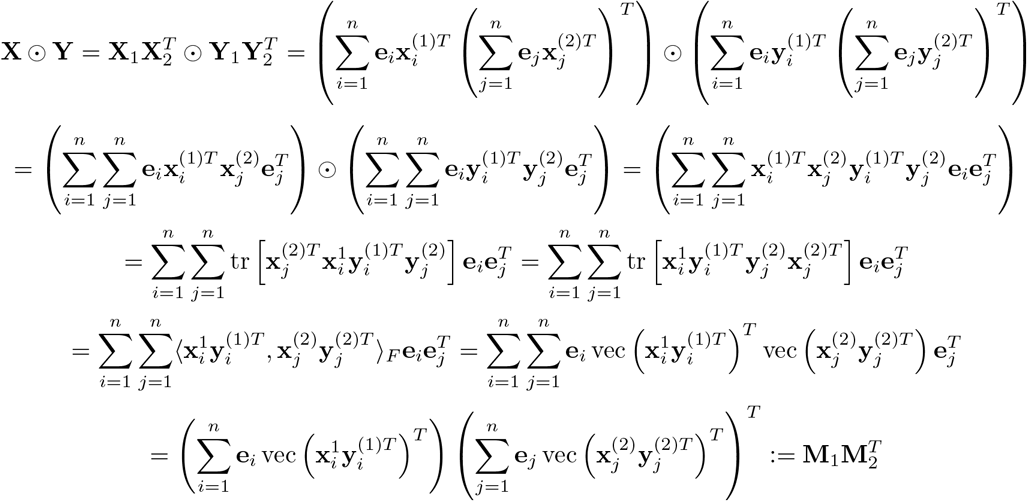

## S5 Clustering from HM-OT output

Suppose we have a time series **X** = **X**^1^, …, **X**^*N*^, and `HM`-`OT` output: a temporal clustering **Q,T**

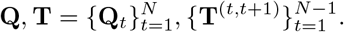

### S5.1 Max-likelihood co-clustering from HM-OT output

The simplest way to extract a hard clustering of each timepoint using **Q,T** is through the **Q**_*t*_ matrices,each of which is *n*_*t*_ × *r*_*t*_ in shape. For each *t* = 1, …, *N*,assign each spot *i* ∈ [*n*_*t*_] to a cluster label 1, …,*r*_*t*_ by taking the argmax over each row. For each *t* ∈ [*N*],this corresponds to clustering function

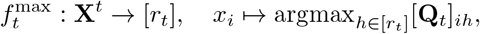

and clustering 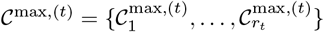 given by

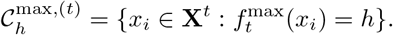

This hard clustering uses the joint probabilities 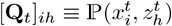.We call this *max-likelihood clustering* of **X** using **Q,T**.

It may also be of interest to instead use the corresponding matrices **Λ**_*t*_ of conditional probabilities for the same purpose. Recall that

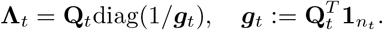

From these, clustering function

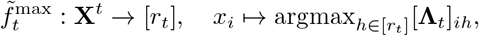

and corresponding clustering 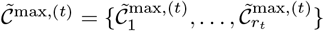 given by

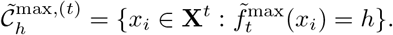

As this hard clustering uses the conditional probabilities 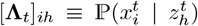,which play the role of emission probabilities in the HMM interpretation of **Q,T**,we call this *emission max-likelihood* clustering of **X** using **Q,T**.

### S5.2 Reference joint clustering from HM-OT output

There is an alternative clustering making use of the latent coupling matrices **T** and a reference timepoint *t*_*_ ∈ [*N*]. All timepoints are then clustered relative to the set of labels 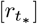 at timepoint *t*_*_.

This method of clustering relies on the fact that,together **Q,T** define a joint distribution between any two timepoints (not necessarily consecutive) that factors through a composition of latent coupling matrices. Without loss of generality,suppose *s,t* ∈ [*N*] with *s* < *t*. Define latent coupling matrix **T**^(*s,t*)^ by

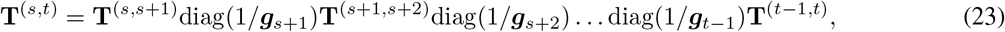

and define the joint distribution of timepoints *s* and *t* from **Q,T** as:

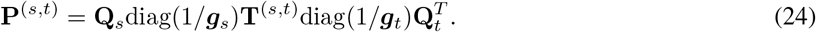

We then define clusterings at each timepoint as follows. Starting with the reference timepoint *t*_*_,we use max-likelihood clustering (described just above) to assign its labels:

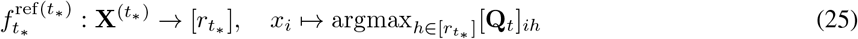

However, for all other timepoints *s*,we use the couplings 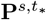 and 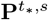 (depending on whether *s* < *t*_*_) to assign cluster labels. Each of these couplings is used to form a map 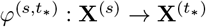, after which labels are assigned to **X**^(*s*)^ by composing with 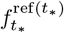 in (25). Concretely,

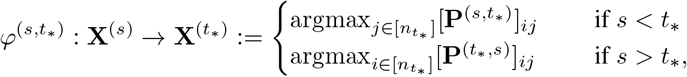

and we define the reference clustering functions at other timepoints *s*:

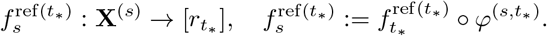

Clustering function 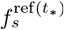 yields a reference clustering of **X**^(*s*)^, relative to timepoint *t*_*_:

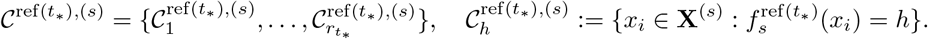

## S6 Metrics

### S6.1 Metrics for Differentiation Maps

#### Pointwise Mutual-Information (PMI) for Differentiation Maps

Another measure of the quality of a differentiation map is the deviation between the probability of a transition between cell types *a* to *b* and the probability of the same transition under the independence model. Specifically,given the joint distribution **T**^(*s,t*)^,defined in 5,and marginal distributions **g**_*s*_,**g**_*t*_ between any pair of timepoints *s,t*,we define the deviation using the pointwise mutual information, an information-theoretic metrics widely used in machine-learning [15]:

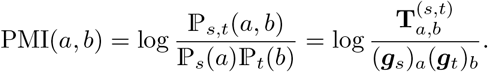

The normalized PMI (NPMI) scales PMI to be in [− 1,+1]: +1 is perfect correspondence,−1 is for never co-occurring,and 0 is for independence.

## S7 Experimental Details

### S7.1 Synthetic Examples

#### S7.1.1 2-timepoint toy model of cell-differentiation

We illustrate the advantages of `HM`-`OT` on a toy model for single-cell differentiation whose dynamics are driven by a “Waddington potential” Φ : ℝ ^*g*^ → ℝ. In the Waddington framework [33,38],the velocity with which cells differentiate depends only on their location in expression-space,descending according to the gradient of the Waddington potential: 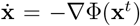,where 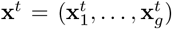 a vector of transcript expression values for some cell at time *t*. For our illustrative example, we simulate cells having *g* = 2 genes,and use a gradient field −∇ Φ defined piecewise in each quadrant of ℝ^2^,described explicitly in Equation 26 and shown in Fig. S1**a**). We evaluate whether `HM`-`OT` can cluster points by quadrant when only given an unlabeled timepoint of the dynamics at a later time. We illustrate the distinction between clusters output by `HM`-`OT`, and those obtained from standard single time-point clustering methods,such as *k*-means. In (Fig. S1**b**),we observe that `HM`-`OT` (i.e. low-rank OT as a sub-case of `HM`-`OT` for 2 time points) correctly identifies the initial condition of the four clusters. Single timepoint clustering methods such as *k*-means are unable to identify such structure and cluster the dataset arbitrarily, only clustering on structure intrinsic to the dataset itself. `HM`-`OT` and low-rank OT are instead able to cluster by an external field (e.g. the Waddington gradient) acting on the data by using temporal information.

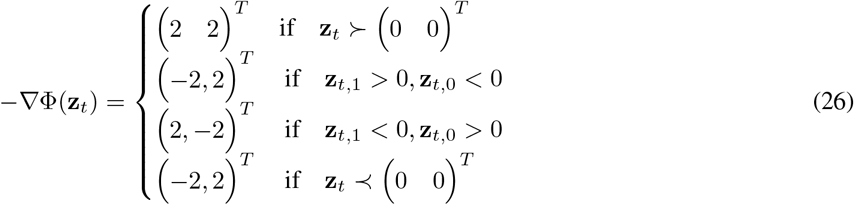

Comparing against *K*-means, one finds the AMI against the ground-truth initial condition for the dynamics, `HM`-`OT` achieves AMI = 0.873 and *K*-means achieves AMI = 0.393. This highlights the advantage of low-rank optimal transport for clustering on the external field a dataset is subject to, as opposed to the intrinsic structure within it alone.

#### S7.1.2 3-timepoint moving Gaussian-mixtures

To demonstrate the value of multiple (≥ 3) timepoints in joint-clustering,we compare a latent pairwise optimal transport method (FRLC) to `HM`-`OT`on a time series dataset with three timepoints *t*_1_,*t*_2_,*t*_3_ (Fig. S1**c**). Timepoint *t*_1_ is sampled from a mixture of two bivariate Gaussians with means 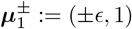;timepoint *t*_2_ is sampled from a single bivariate, centered Gaussian; and timepoint *t*_3_ is sampled from a mixture of two bivariate Gaussians with means 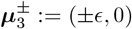.All Gaussians in this example are isotropic, with covariance matrix given by *σ*^2^ times the identity. Pairwise latent optimal transport identifies cost-minimizing clusters at *t*_2_ for each pairwise direction (*t*_1_ →*t*_2_ and *t*_2_ →*t*_3_). The resulting clusters in each case (Fig. S1**d**) have orthogonal boundaries, reflecting the relative positions of *t*_1_ or *t*_3_ to *t*_2_, rather than a consistent differentiation polarity across all three timepoints. `HM`-`OT` identifies clusters minimizing the total transport cost *t*_1_→ *t*_2_ →*t*_3_, and identifies a hyperplane separassssting progenitor-like and descendant-like cells at *t*_2_ using all three time-points. Whereas pairwise optimal transport minimizes according to first-order velocities, `HM`-`OT` captures second order information on trajectories, such as curvature or momentum.

**Figure S1:**
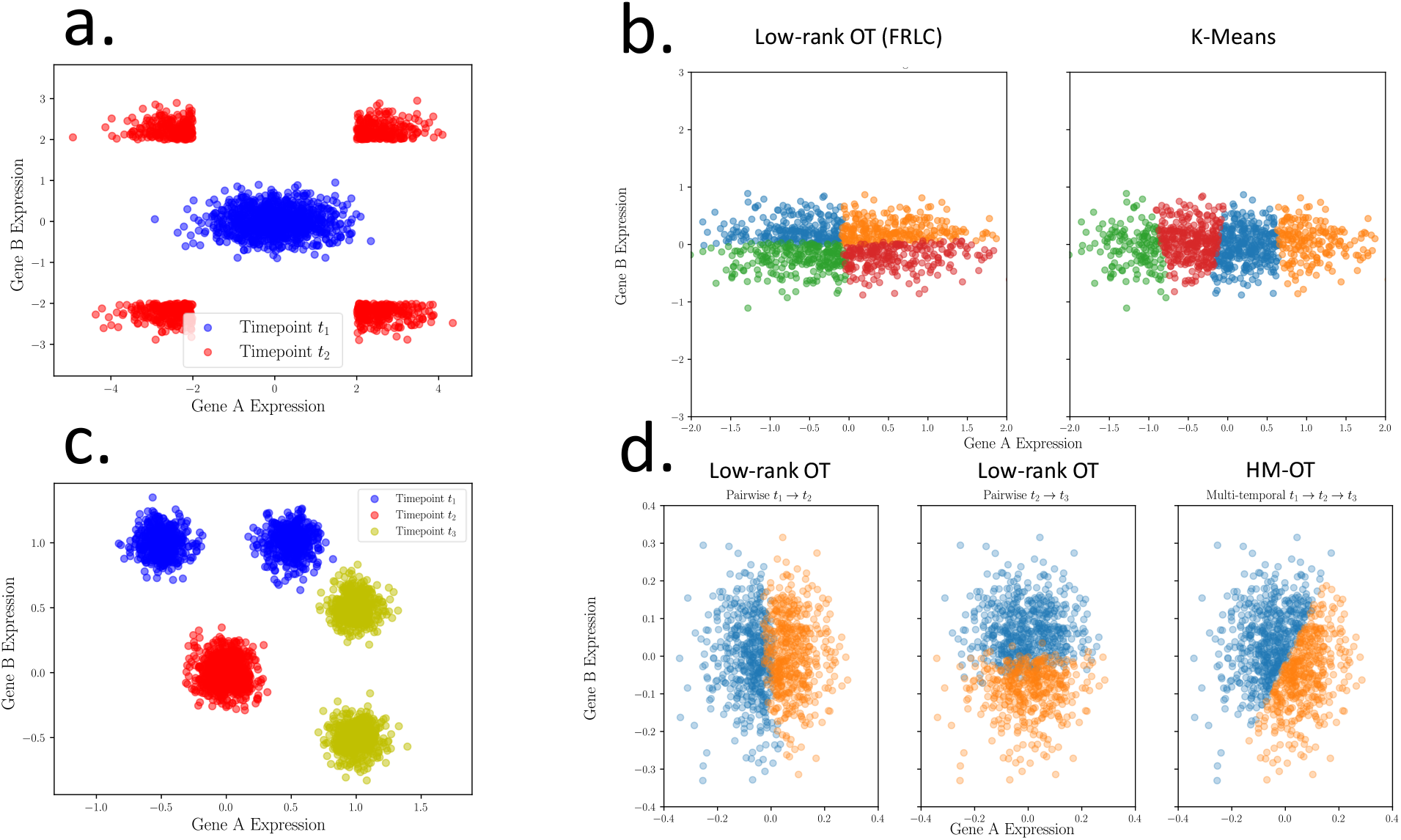
HM-OT on a synthetic example. **a**.An example of data from two timepoints with a vector field splitting on four quadrants – the initial condition of *t*_1_ determines which of the four clusters the cell differentiates into. **b**. Pairwise clustering using `HM`-`OT` (2 timepoints),and clustering using *k*-means. **c**. A three-timepoint synthetic example colored according to clusters which minimize respective pairwise or ternary losses. **d**. (Left) Clustering at time *t*_2_ obtained by a pairwise alignment of *t*_1_ and *t*_2_; (Middle) The clustering at time *t*_2_ obtained by a pairwise alignment of *t*_2_ and *t*_3_; (Right) Clustering at *t*_2_ obtained via `HM`-`OT` alignment of all three timepoints.

### S7.2 Zebrafish embryogenesis, spatial dataset

#### S7.2.1 Preprocessing and Hyperparameter Selection

We demonstrate the extension of `HM`-`OT` to Zebrafish spatial transcriptomics, using the Stereo-Seq dataset of [21] across 3.3hpf, 5.25hpf, 10hpf, 12hpf, 18hpf, and 24hpf (where hpf denotes hours post fertilization). We use `scanpy` to load the datasets. To generate pairwise costs we take the intersection of common genes between each pair of timepoints to generate a joint AnnData object. Given this joint AnnData, containing all transcriptomic features at both timepoints, we perform a normalization with `scanpy.pp.normalize_total` and add log pseudocounts with `scanpy.pp.log1p`. We project the transcript vectors onto the first 60 principle components of the joint dataset **UU**^*T*^ **X**^(*i*)^, computed using `scanpy.pp.pca`. The pairwise costs **C**^(*i,i*+1)^ are computed using these components. In Table S3 we list the hyperparameters used on the spatial Zebrafish dataset. *α* > 0 implies we use a spatial Gromov-Wasserstein term in the objective, to distinguish it from the Wasserstein-only (single-cell) case. We use a Geodesic cost matrix for this GW-term and factorize it with an SVD, as the method takes a distance matrix as input in low-rank form.

#### S7.2.2 Differential Expression Analysis for Notochord and Adaxial

The somite and adaxial cell types are approximately divided between `HM`-`OT` types (13) (*N* = 333 spots) and (15) (*N* = 395 spots), where (13) is predicted by `HM`-`OT` to transition to notochord and (15) is predicted to transition to somite (Fig. 2**a**). Somites are mesodermal pre-muscular cells, and adaxial cells differentiate within the somites to muscle fibers. The notochord, meanwhile, is a rod-like structure which supports the neural tube and is surrounded by adaxial cells [24, 5]. Interestingly, the (13) cell type is more central and flanked by type (15) on both sides–recapitulating how the notochord is a central rod flanked by both somite and adaxial cells [24, 5]. We perform differential expression using a T-test with Benjamini-Hochberg correction. The “adaxial” cell type of [21] has top 5 genes *acta1a* (*Z* = 20.68, for *α*-actin in skeletal muscle), *myl10* (*Z* = 19.69, a myosin muscle protein), *hspb1* and *hsp90aa1*.*1* (*Z* = 19.45 and = 11.26, both heat shock factors), and *chrd* (*Z* = 10.87). Our cell type (15) has 4 of the same genes among the top 5 genes *acta1a* (*Z* = 10.61), *myl10* (*Z* = 10.17), *hspb1* (*Z* = 9.96), *hsp90aa1*.*1* (*Z* = 8.30), but has *ripply1* (*Z* = 6.60) as the fifth most represented instead of *chrd*, which is no longer among even the top 10 differentially expressed in cell type (15). Chordin is a key notochord marker, and *ripply1* is a gene involved in somite segmentation which also is expressed in the notochord type [16]. None of the top 10 genes in this identified type are muscular, with the expression of the skeletal muscle proteins in adaxial substantially diminished with *acta1a* (*Z* = 4.53) and *myl10* (*Z* = 4.76) substantially less expressed. This recapitulates the established co-localization of the distinct adaxial-somite and notochord cell types [24, 5].

#### S7.2.3 Further discussion of NPMI values between 10hpf and 18hpf

Both the `HM`-`OT` cell types and [21] suggest yolk-syncytial has NPMI = 0.312 through the graph. Both maps erroneously suggest differentiation occurring from the yolk-syncytial layer – this is partially due to the balanced OT constraint which requires all the cells of yolk-syncytial transfer, and perhaps error in the annotation. It is established that ectodermal cells transition to the neural plate and then the nervous system and neural rod [19]. `HM`-`OT` suggests an ectodermal layer of cells, labeled “periderm,” gives rise a neural plate intermediate which differentiates into the nervous system. The transitions from `HM`-`OT` and annotation [21] cell types both suggest that the neural keel, differentiates into a neural plate intermediate and transitions in a second trajectory to neural crest. The neural keel and neural crest are differentiated from the neural plate, and the transition of neural plate to neural crest is supported in the literature [12]. Observing Figure 2c and d, we see cell types (7) and (9) are consistent with the segmentation of the 18hpf neural crest and nervous system in a manner not captured by the 12hpf annotations.

**Table S1:** NPMI values of HM-OT cell types on developmental zebrafish dataset of [21].

|  | Anterior Neural Keel | Notochord | Paraxial Mesoderm, Neural Keel | Periderm | Posterior Neural Keel | Segmental Plate, Tail Bud | Yolk Syncytial Layer |
| --- | --- | --- | --- | --- | --- | --- | --- |
| Angioblastic Mesenchymal Cell | -0.917 | -0.943 | -0.905 | -0.921 | -0.891 | -0.969 | 0.517 |
| Erythroid Lineage Cell | -0.973 | -0.942 | -0.976 | -0.979 | -0.935 | 0.52 | -0.969 |
| Forebrain | -0.935 | -0.96 | -0.868 | -0.089 | -0.919 | -0.977 | 0.618 |
| Hatching Gland | -0.991 | -1.0 | -1.0 | -0.992 | -1.0 | -1.0 | 0.488 |
| Immature Eye, Midbrain | -0.93 | -0.95 | -0.882 | 0.23 | -0.919 | -0.967 | 0.194 |
| Nervous System | 0.454 | 0.011 | -0.147 | 0.517 | 0.288 | -0.926 | -0.898 |
| Neural Crest | 0.17 | -0.737 | 0.813 | 0.174 | 0.114 | -0.954 | -0.775 |
| Neural Crest, Otic Vesicle | 0.37 | -0.611 | -0.081 | 0.402 | 0.262 | -0.924 | -0.859 |
| Notochord | -0.966 | 0.665 | -0.969 | -0.976 | -0.857 | 0.033 | -0.985 |
| Pronephros | -0.939 | -0.82 | -0.947 | -0.944 | -0.886 | 0.213 | -0.96 |
| Segmental Plate, Tail Bud | -0.702 | 0.252 | -0.749 | -0.722 | -0.651 | -0.032 | -0.892 |
| Somite | -0.96 | -0.264 | -0.962 | -0.975 | -0.837 | 0.497 | -0.986 |
| Yolk Syncytial Layer | -0.989 | -1.0 | -1.0 | -0.99 | -1.0 | -1.0 | 0.312 |
| Yolk Syncytial Layer, Periderm | -0.985 | -0.987 | -0.987 | -0.986 | -0.981 | -0.984 | 0.38 |

**Table S2:**
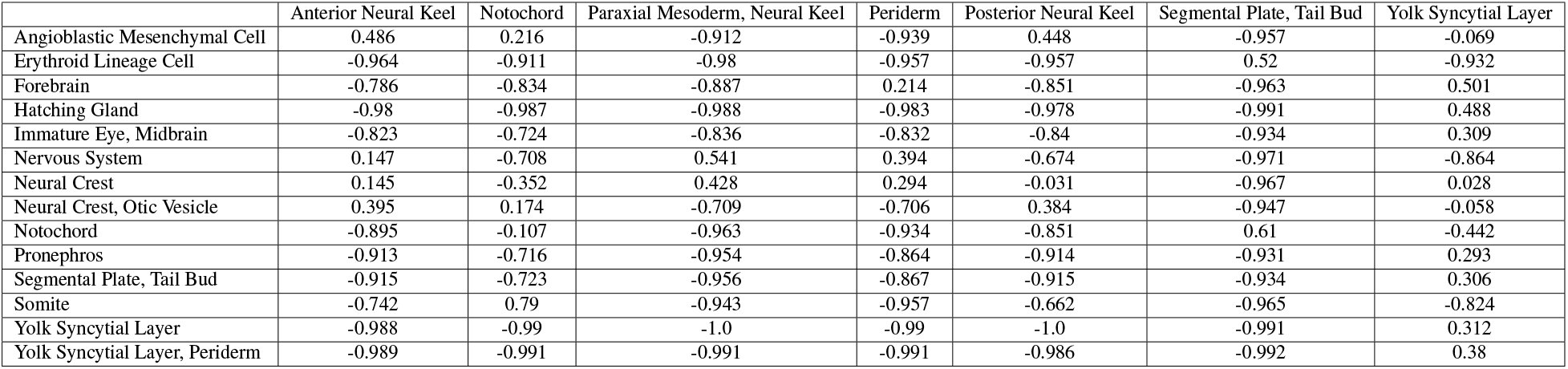
NPMI values of annotated cell types on developmental zebrafish dataset of [21].

#### S7.2.4 Validation of unsupervised differentiation map in minimizing transcriptional distance to *unseen* costs

Using `HM`-`OT` we infer the differentiation map either with the annotated clusters or the unsupervised clusters `HM`-`OT` can additionally learn. To validate the biological accuracy of these unsupervised clusters and their differentiation map, we compute pairwise transcriptional distances between stages separated by greater than one timepoint, `HM`-`OT` has no access to as it only uses pairwise distances between adjacent timepoints. A reasonable differentiation map, under compositions of its different stages, should align ancestral and descendant clusters at time *i* and those at j for j > *i* + 1 with least transcriptional distance. This is demonstrated in Figure **??**, where we show that using the unsupervised clusters the composition of the differentiation map between all timepoints up to 24hpf has lower average transcriptional distance than the differentiation map using annotated clusters: the improvement in the TD ranges from an average of 6.24 transcripts/spot at 18hpf to near zero at 3hpf, which is fully undifferentiated and distant from all differentiated 24hpf clusters.

**Table S3:** HM-OT model Hyperparameters.

| Hyperparameter | Description | Value |
| --- | --- | --- |
| $\alpha; (1 - \alpha)$ | Weight to Gromov-Wasserstein; to Wasserstein | 0.01; 0.99 |
| $\tau_{out}$ | Regularization of outer marginals deviating from $\mathbf{a}$ or $\mathbf{b}$ | 1 |
| $\tau_{in}$ | Mirror-descent regularization of inner marginal at step $g_Q^k$ deviating from $g_Q^{k-1}$ | $1 \times 10^{-7}$ |
| $\gamma$ | Mirror-descent step-size | 30 |

**Table S4:** Hyperparameters of `HM`-`OT` on developmental zebrafish dataset of [21].

#### S7.2.5 Unsupervised Cluster Loss on Standard OT-Costs

As a simple baseline, we also evaluate whether the inferred unsupervised clusters from Problem 2, where (**Q,T**) are jointly minimized, achieve a lower value of the Wasserstein and Gromov-Wasserstein loss than the supervised clusters from Problem 3 where **T** is minimized and **Q** is fixed from annotation [21]. We demonstrate, for a Fused GW objective in Table S5 that, unsurprisingly, the cost is lower when **Q** is also learned.

**Table S5:**
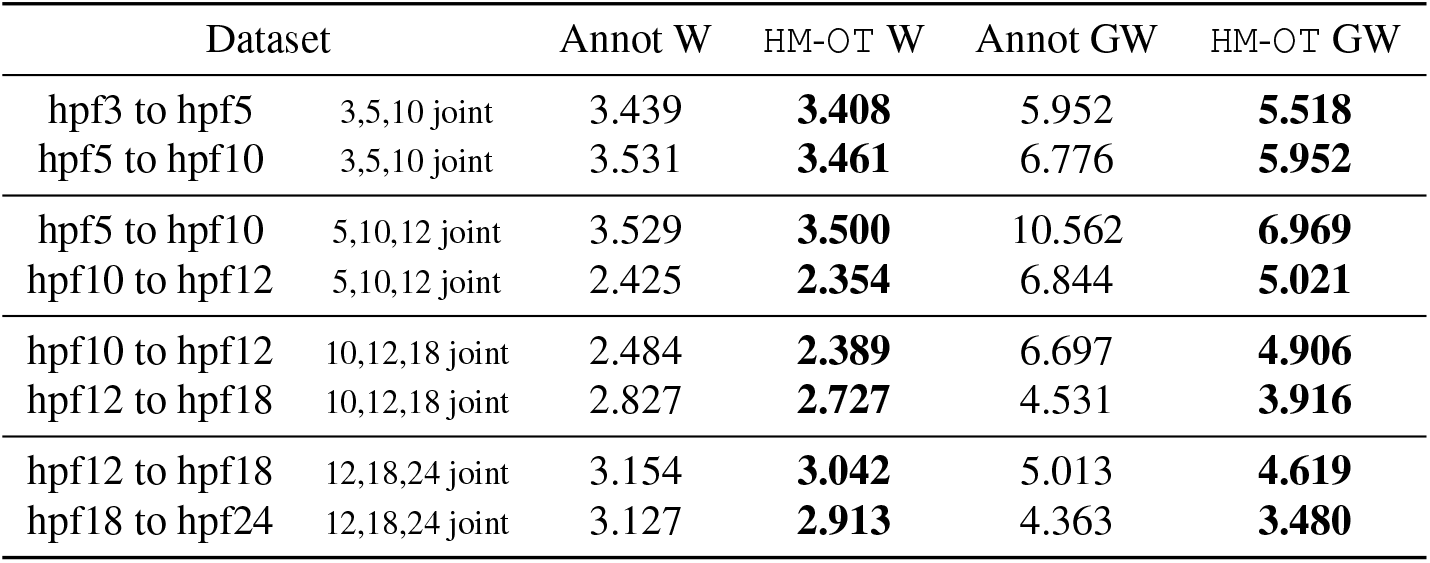
Performance of HM-OT clusters (Algorithm 1) against annotation clusters in minimizing the Wasserstein and Gromov-Wasserstein distance across four joint alignments of time triples (*t*_1_, *t*_2_, *t*_3_). For each clustering, the optimal transport cost for the associated clusters is computed and shown – for this spatial dataset, a larger contribution has been given to the GW-term.

### S7.3 Mouse Embryonic development

#### S7.3.1 Preprocessing and Hyperparameter Selection

We follow the exact procedure for preprocessing outlined in S7.2.1, and offer the hyperparameters used for generating the mouse embryo differentiation map (Figure 3) in S7.

#### S7.3.2 Evaluating Scalability

To evaluate scalability, we ran `HM`-`OT` on all time pairs of the MOSTA Stereo-Seq dataset [3]. We evaluate the performance on low-rank pairwise alignment (setting `max_iter=60`, as used in the experiments) for direct comparison with `moscot`. Unlike `moscot`, `HM`-`OT` can run on all timepoints of the dataset, up to E16.5 which has *N* = 121767 spots. Moreover, the scaling of runtime with dataset size is minimal (Figure S8a) – this is due to the FRLC algorithm which scales low-rank OT using Sinkhorn [4], as opposed to previous works which relied on the less efficient Dykstra subroutine [30, 8, 20] which is used in `moscot` [17]. We also show the improvement in space complexity of `HM`-`OT` in using a low-rank coupling factorization over storing the quadratic full-rank matrix (Figure S8b).

**Table S6:** Runtimes of moscot and HM-OT across different time-points in seconds.

|  | E9.5_10.5 | E10.5_11.5 | E11.5_12.5 | E12.5_13.5 | E13.5_14.5 | E14.5_15.5 | E15.5_16.5 |
| --- | --- | --- | --- | --- | --- | --- | --- |
| `moscot`<br>(solve only) | 13.950 | 25.647 | 24.347 | N/A | N/A | N/A | N/A |
| HM-OT | 8.985 | 9.077 | 9.105 | 9.141 | 9.251 | 9.304 | 9.311 |

**Table S7:** Hyperparameters of HM-OT on MOSTA [3] Stereo-Seq mouse organogenesis dataset.

| Hyperparameter | Description | Value |
| --- | --- | --- |
| $\alpha; (1 - \alpha)$ | Weight to Gromov-Wasserstein; to Wasserstein | $10^{-4}; 1 - 10^{-4}$ |
| $\tau_{out}$ | Regularization of outer marginals deviating from $\mathbf{a}$ or $\mathbf{b}$ | 5 |
| $\tau_{in}$ | Mirror-descent regularization of inner marginal at step $\mathbf{g}_Q^k$ deviating from $\mathbf{g}_Q^{k-1}$ | $1 \times 10^{-5}$ |
| $\gamma$ | Mirror-descent step-size | 40 |

## S8 Supplemental Figures

**Figure S2:**
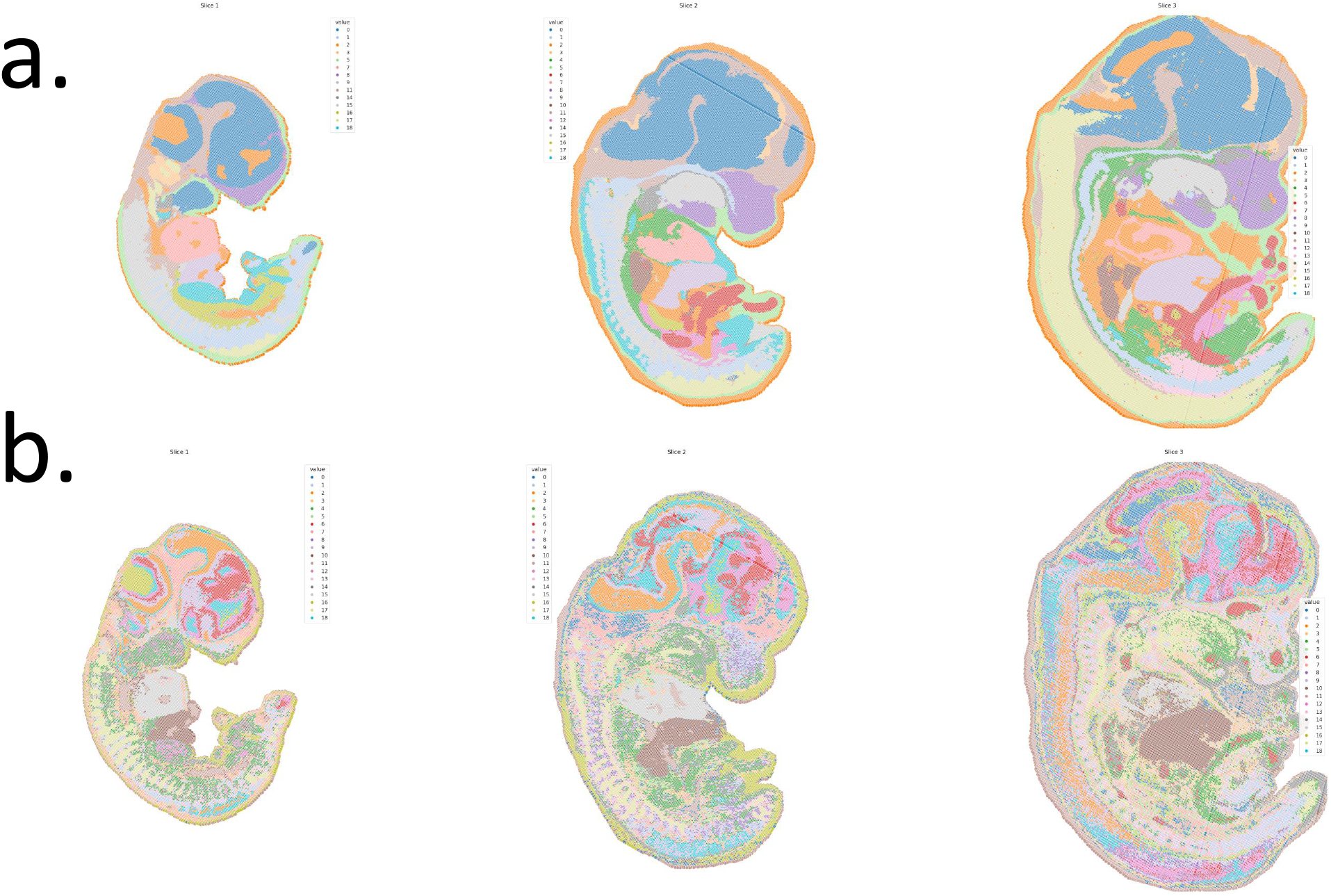
Ancestral clustering on the MOSTA mouse organogenesis dataset of [3] with reference index 5 on **a**. the [3] annotations and the `HM`-`OT` differentiation map for linking them, **b**. `HM`-`OT` unsupervised cell types. As the reference index is the final timepoint, this corresponds to “pulling back” differentiated cell types to their progenitors.

**Figure S3:**
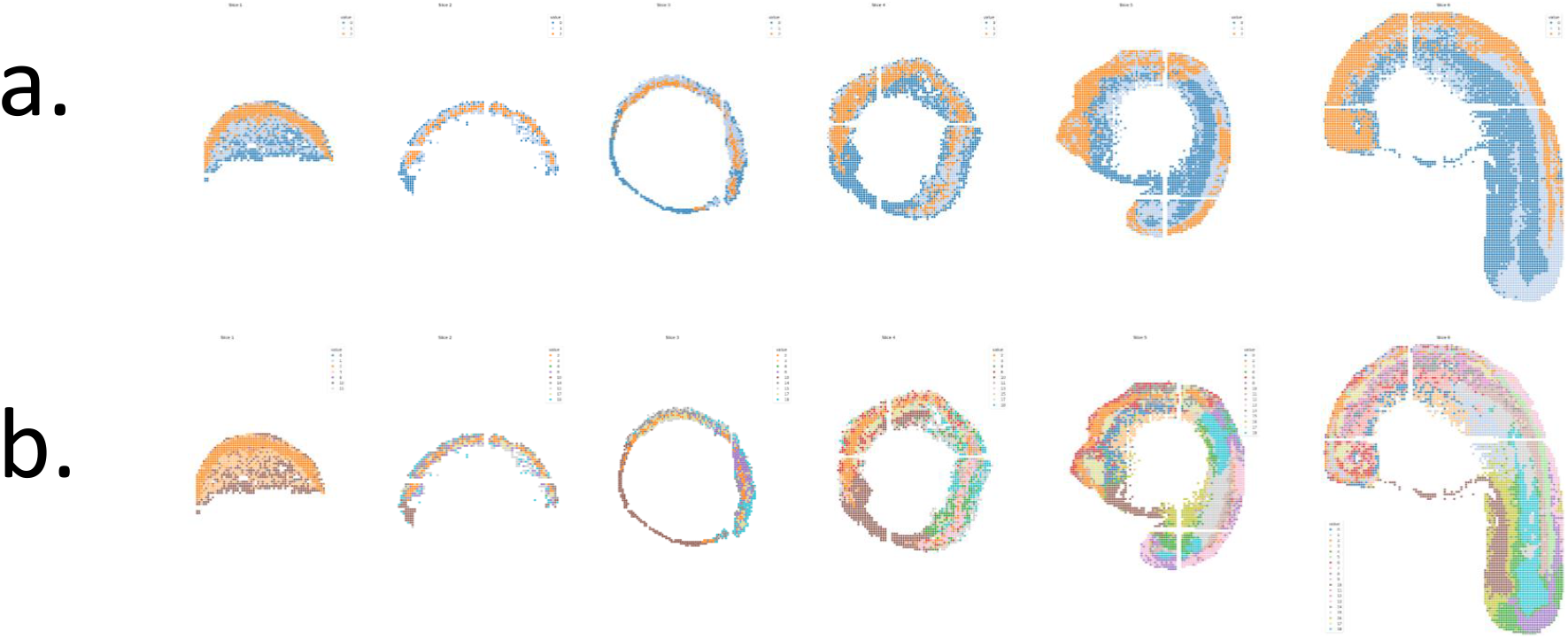
Ancestral clustering on the zebrafish dataset of [21] with fully-unsupervised cell types inferred by `HM`-`OT` with **a**. reference index zero corresponding to mapping progenitor cells into future descendant locations, **b**. ancestral clustering with reference index 5, corresponding to “pulling back” differentiated cell types to their progenitors.

**Figure S4:**
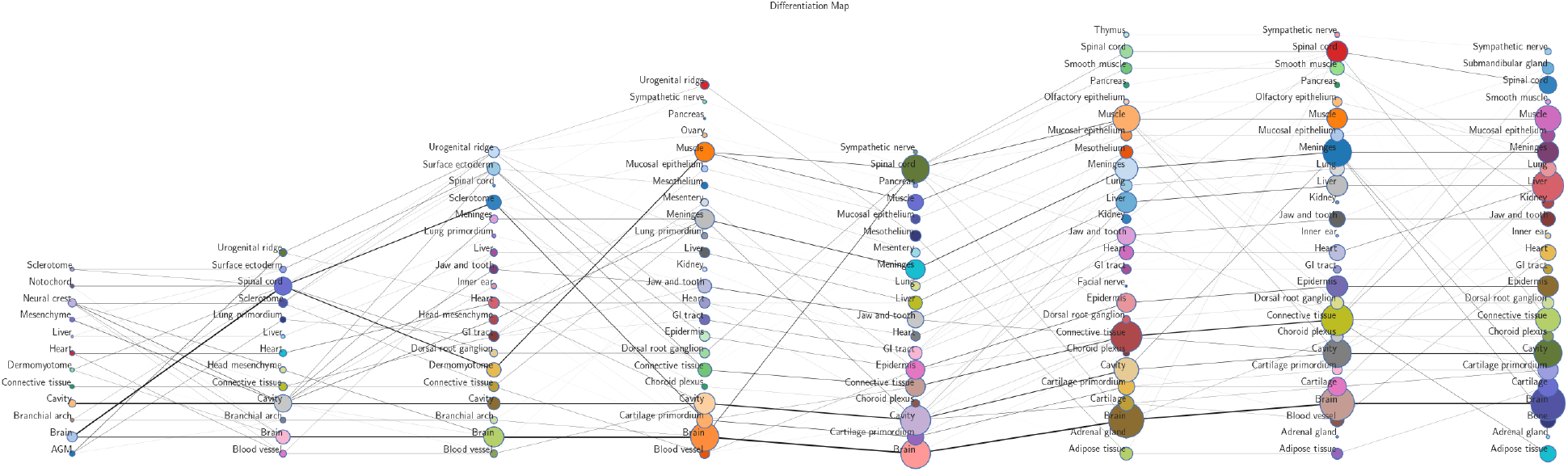
Full differentiation map of Stereo-Seq mouse organogenesis (MOSTA) dataset [3] from timepoints E9.5-E16.5 generated using `HM`-`OT`.

**Figure S5:**
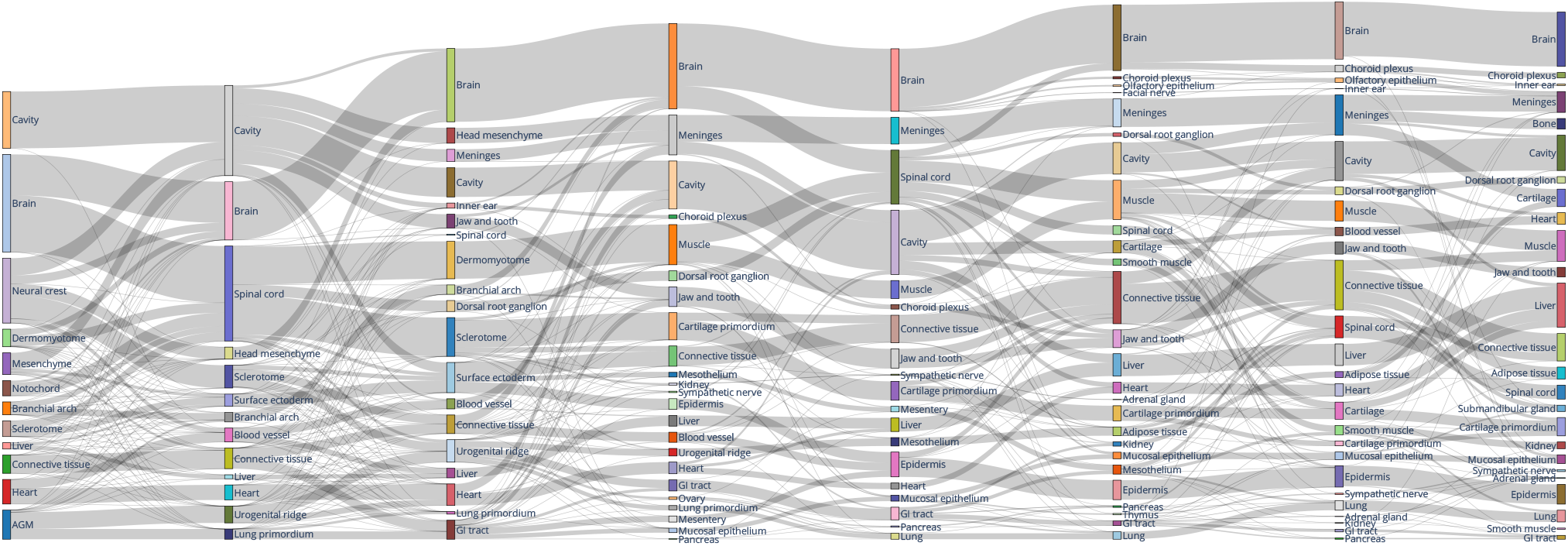
A Sankey plot corresponding to the differentiation map in Figure S4, from E9.5 to E16.5.

**Figure S6:**
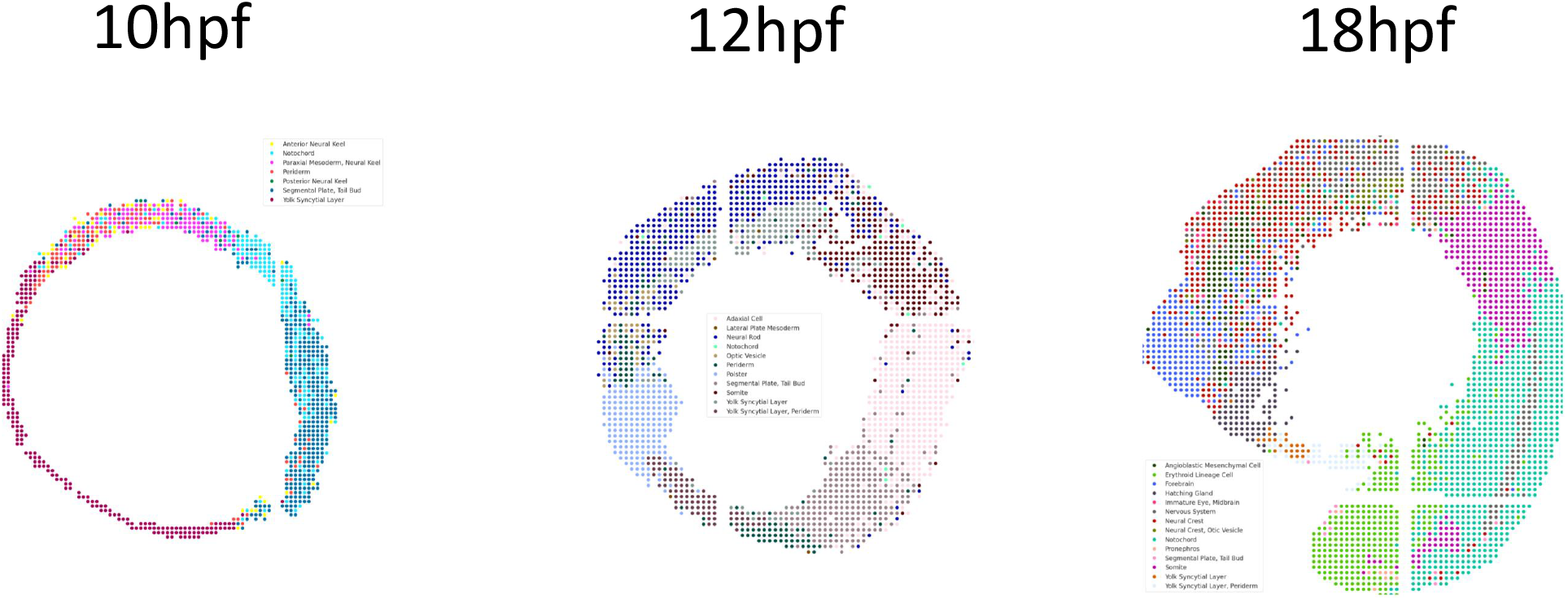
Annotated cell types of [21] plotted spatially for the 10hpf, 12hpf, and 18hpf timepoints.

**Figure S7:**
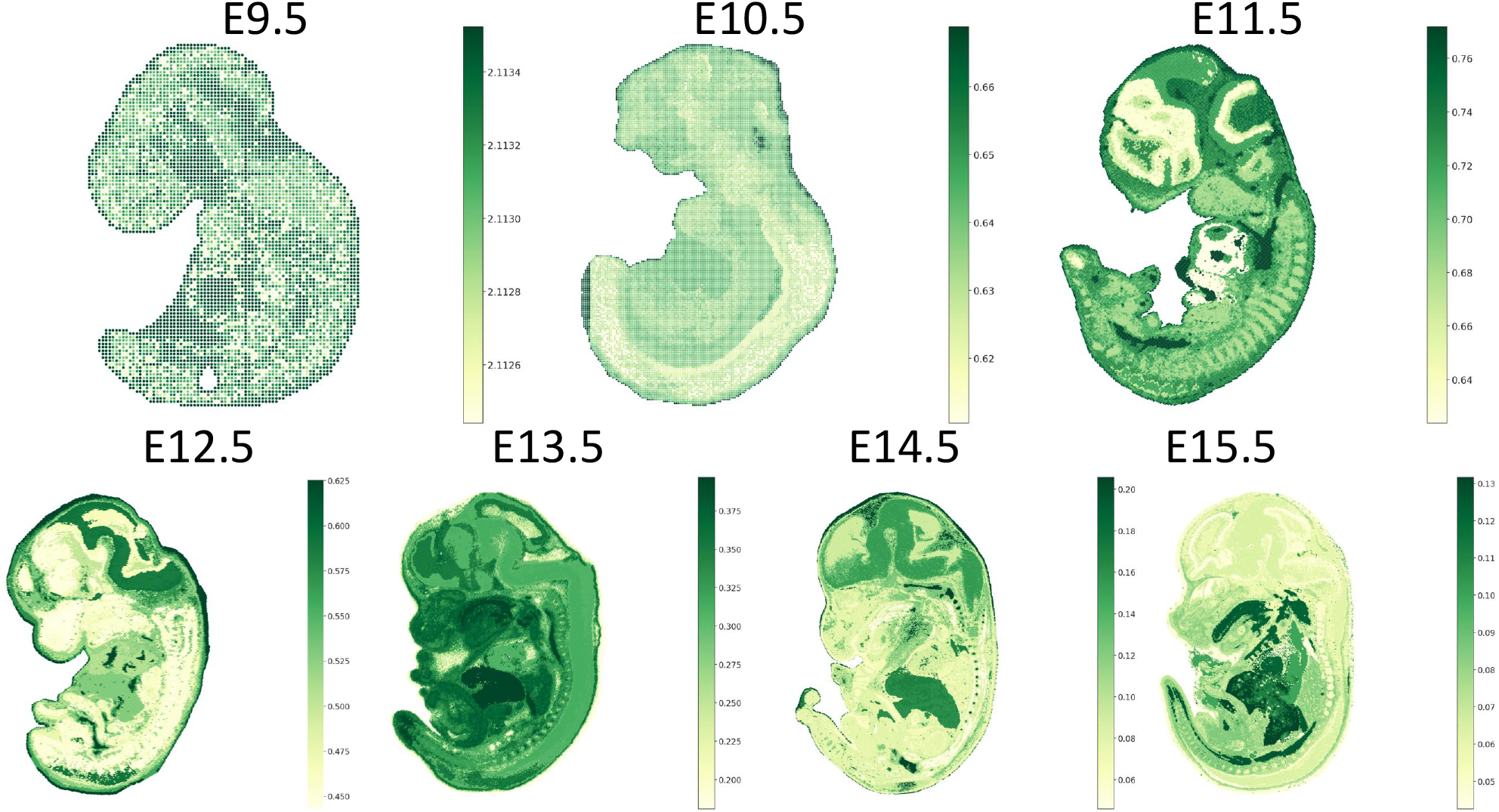
Growth-rates on of Stereo-Seq mouse organogenesis (MOSTA) dataset [3] growth-rates (***ξ***) on stages E9.5-15.5 computed in accordance with [10] on the unsupervised `HM`-`OT` clusters. Growth-rates are scaled to represent the expected number of cells added and sub-couplings computed scalably using the FRLC low-rank optimal transport framework [11].

**Figure S8:**
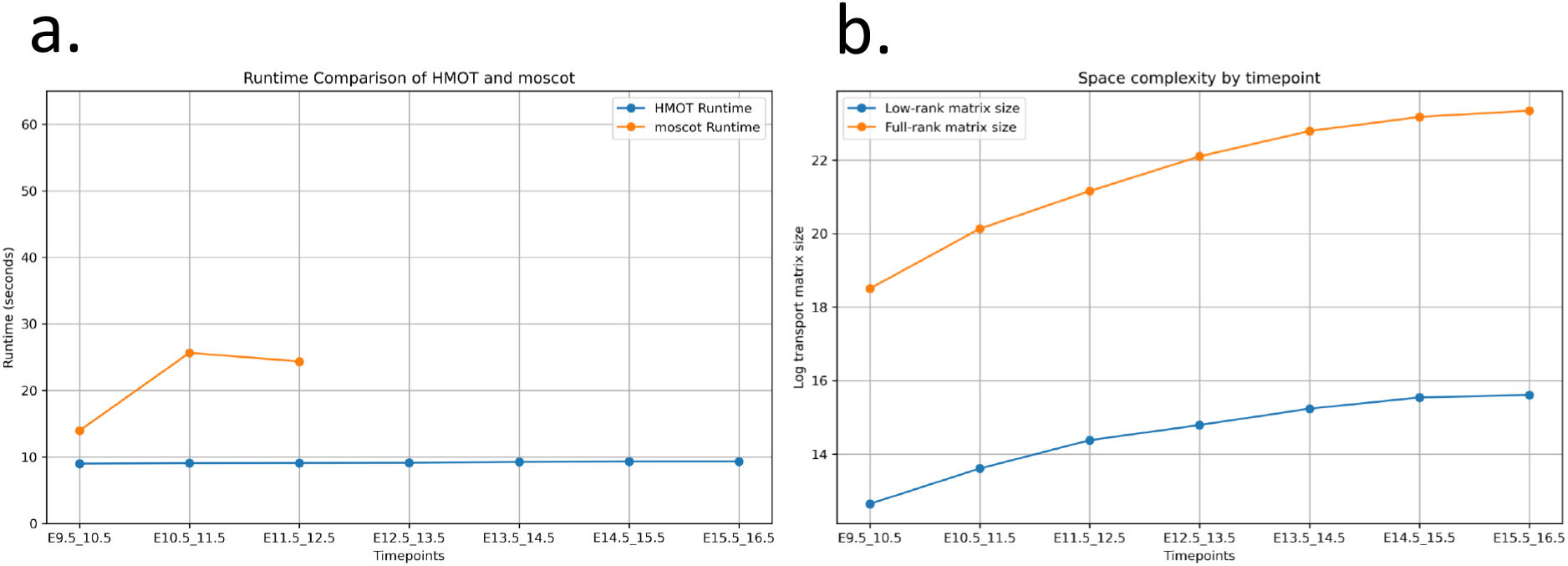
**a**. The runtime of `moscot` (`SpatioTemporalProblem.solve` runtime only) and `HM`-`OT` across pairwise alignments from E9.5-10.5 to E15.5-16.5. `moscot` fails to run after E11.5-12.5. **b**. The memory-scaling of the low-rank transport matrix solved for by `HM`-`OT` versus the analogously shaped full-rank matrix.

## Footnotes

1 *i*_*t*_ denotes an index into the point in **X**^*t*^ on the trajectory, and *k*_*t*_ denotes an index into the set of states 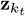 can take

